# Optimizing experimental design for genome sequencing and assembly with Oxford Nanopore Technologies

**DOI:** 10.1101/2020.05.05.079327

**Authors:** John M. Sutton, Joshua D. Millwood, A. Case McCormack, Janna L. Fierst

## Abstract

**Background:** High quality reference genome sequences are the core of modern genomics. Oxford Nanopore Technologies (ONT) produces inexpensive DNA sequences in excess of 100,000 nucleotides but high error rates make sequence assembly and analysis a non-trivial problem as genome size and complexity increases. To date there has been no comprehensive attempt to generate robust experimental design for ONT genome sequencing and assembly. In this study, we simulate ONT and Illumina DNA sequence reads for the model organisms *Escherichia coli*, *Caenorhabditis elegans*, *Arabidopsis thaliana,* and *Drosophila melanogaster* and assemble with Canu, Flye, and MaSuRCA software to quantify the influence of sequencing coverage and assembly approach. Heterozygosity in outbred eukaryotes is a common problem for genome assembly. We show broad applicability of our methods using real ONT data generated for four strains of the highly heterozygous nematode *Caenorhabditis remanei* and *C. latens*.

**Results:**

- ONT libraries have a unique error structure and high sequence depth is necessary to assemble contiguous genome sequences.
- As sequence depth increases errors accumulate and assembly statistics plateau.
- High-quality assembled sequences require a combination of experimental techniques that increase sequence read length and computational protocols that reduce error through correction, read selection and ‘polishing’ with higher accuracy short sequence reads.
- Our robust experimental design results in highly contiguous and accurate genome assemblies for the four strains of *C. remanei* and *C. latens*.

**Conclusions:** ONT sequencing is inexpensive and accessible but the technology’s error structure requires robust experimental design. Our quantitative results will be helpful for a broad array of researchers seeking guidance for *de novo* assembly projects.

## Background

Many factors affect the quality of a *de novo* genome assembly. Genome size increases the size of the “puzzle” to put together, while the size of the pieces (sequence reads) remains the same. Repetitive regions and mobile genetic elements create unique challenges as they are present in more than one location in the genome and without contextual information it is difficult to identify how many copies exist in the complete genome. For example, *Alu* repeat elements reach >1 million copies in the human genome [1].

Third generation sequencing technologies can theoretically solve these problems. The long sequence reads span repetitive regions, potentially allowing for the identification of the exact size and location of repeats on a chromosome. Long sequence reads increase the puzzle piece size for assembly and require less sequencing effort to span the entire genome. Pacific Biosciences (PacBio) and Oxford Nanopore Technologies (ONT) are the current front-runners in long-read sequencing platforms; both are capable of average read lengths in the tens of thousands of base pairs and, theoretically, entire chromosomes can be sequenced in a single read [2, 3]. Both are also capable of producing quality genome assemblies with reasonable amounts of data [4].

ONT offers several advantages over PacBio. Nanopore sequencing relies on running molecular fragments through engineered nanopores and recording the resulting alterations in electrical current. The technology is versatile and can be used for DNA sequencing, mRNA sequencing, amplification-free mRNA quantification [5], and measuring DNA base modifications like methylation [6, 7]. ONT libraries can be readily prepared with low amounts of input DNA, which is an important consideration when studying organisms that are small or difficult to sample. ONT platforms are inexpensive and designed to be used in individual research laboratories and classrooms [8]. For these reasons we have chosen to study ONT and quantify how this inexpensive, accessible technology may be best utilized to produce high-quality assembled reference genome sequences. Similar projects analyzed experimental design for PacBio [9, 10] but the few benchmarking studies for ONT sequencing and assembly are carried out on a narrow scope of organisms [11–13]

ONT sequence reads present unique challenges for genome sequence assembly. DNA molecules do not move through protein nanopores at a constant rate and changes in current are a composite signature reflecting 3-5 nucleotides occupying the nanopore (for R9.4.1 flow cells). The signal processing has trouble detecting changes in current with homopolymers (single nucleotide repeats, for example AAAAAA), short tandem repeats and heavily methylated sites [14, 15]. As a result, ONT sequence reads contain small nucleotide stretches that have been incorrectly identified, inserted and deleted (Fig. 1A, C). This error structure results in relatively few large, contiguous stretches of correctly identified nucleotides (Fig. 1B, D) and is uniquely challenging for assembly algorithms.

**Figure 1.**
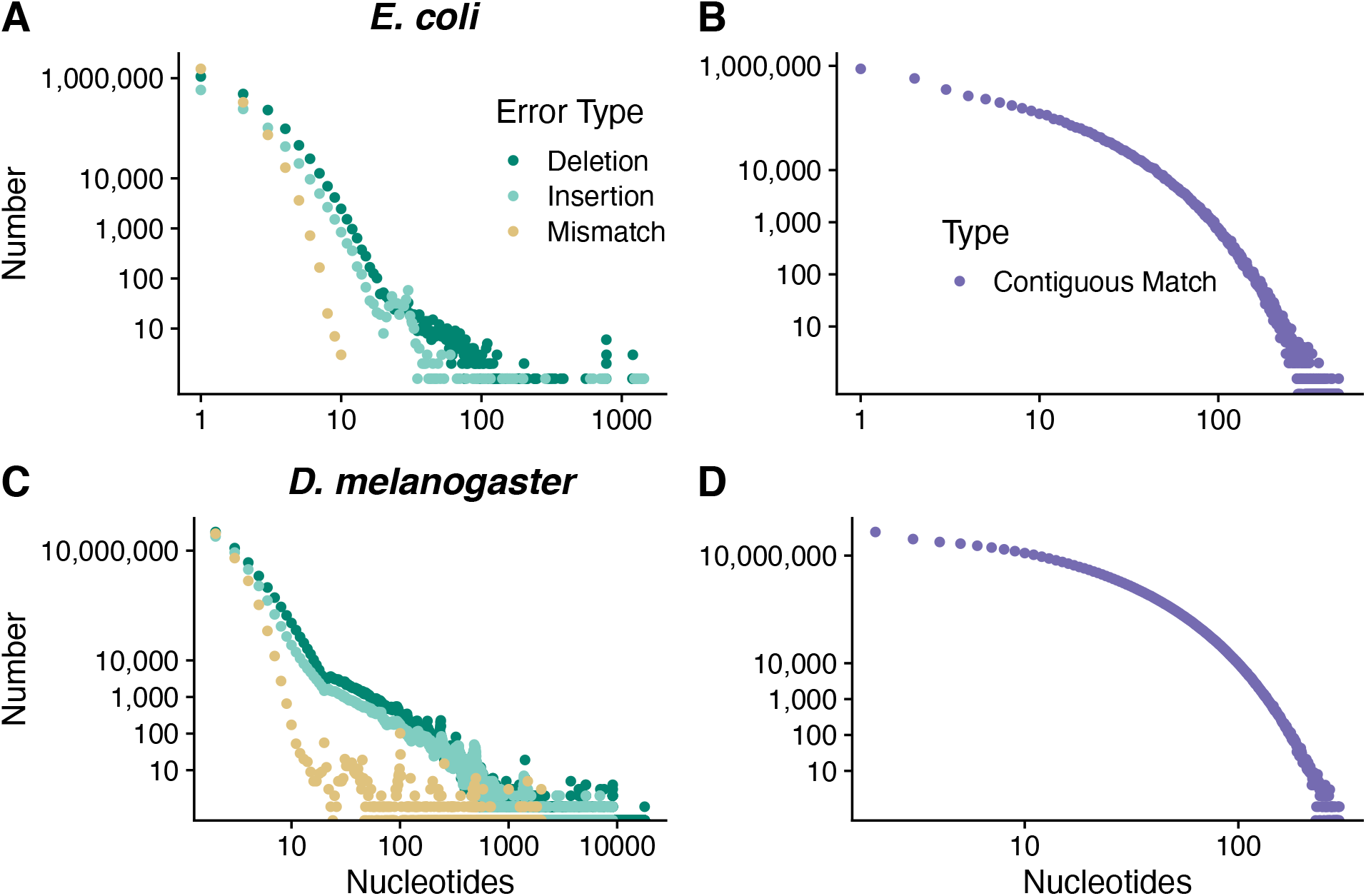
ONT sequence reads contain mixtures of errors including miscalled nucleotides, deletions, insertions and truncated homopolymers. When aligned to the reference genome this results in a large number of single and multi-base deletions, insertions and mismatches for (A) *E. coli* and (B) relatively few stretches of contiguous matching sequence that extend beyond a few nucleotides. For (C) *D. melanogaster* the error profile is similar but the large, repeat-rich genome results in multi-nucleotide deletions and insertions and again (D) few stretches of long, contiguous matching sequence.

To achieve our results we simulate ONT DNA libraries for the model organisms *Escherichia coli*, *Caenorhabditis elegans*, *Arabidopsis thaliana,* and *Drosophila melanogaster*. For each organism we assemble sequence read sets at different depths (i.e., the average times a nucleotide in the genome is sequenced). We measure the contiguity, completeness, and accuracy of each assembled sequence relative to the current reference genome. Many *de novo* assembly projects target organisms without reference genomes and we also measure the identification of a set of genes thought to be found in single copy in each organism. Pure ONT datasets result in superior assembled genome sequences but can be cost-prohibitive for large genomes where multiple flow cells may be necessary to gather enough data. We also analyze assembled sequences from ‘hybrid’ DNA read datasets composed of ONT and Illumina sequence reads.

For eukaryotes both ploidy and breeding system can influence the assembly of genome sequence. Polyploidy can create scenarios where many sites in a genome look highly similar to one another, making it tough to place these regions within an assembled genome [16, 17]. For non-haploid organisms, there is a potential for heterozygosity and at these sites the effective sequencing coverage is cut in half. For small, non-clonal metazoans sufficient DNA for sequencing cannot be acquired from a single individual. DNA must be extracted from pools of individuals, and for outbreeding organisms there may be substantial individual diversity within the pool. Here, we use results from our simulated sequences to guide experimental ONT sequencing and assembly for four strains of the highly heterozygous *Caenorhabditis remanei* and *C. latens*. We present the experimental and computational protocols that result in high quality assembled genome sequences.

We find that ONT approaches can produce contiguous assembled sequences for the organisms we study with relatively high sequencing coverage of >60x, well above current recommendations [18]. Pure ONT sequencing and assembly outperforms our tested hybrid approach. We find that contiguous assemblies cannot be achieved by solely increasing ONT sequencing depth as errors accumulate and assembly statistics plateau. Pre-assembly filtering and read correction improve contiguity, and post-assembly polishing using short Illumina DNA sequence reads increases accuracy. We also find that the use of Illumina data, even at very low sequencing depths, increases accuracy through iterative polishing. Our results demonstrate that rigorous experimental design can improve our ability to infer a genome sequence and reduce the effort and cost required.

## Data Description

### Simulated sequence libraries

We obtained ONT sequences (R9 chemistry) from the National Center for Biotechnology Information (NCBI) Sequence Read Archive (SRA) on February 2, 2019 [19]. The *E. coli* dataset contained 11,652,330 sequenced bases in 120,151 reads (~2.5x coverage of the 4.64Mb reference sequence), the *C. elegans* dataset contained 8,860,671,330 sequenced bases in 583,466 reads (~87.9x coverage of the 100.8Mb reference sequence), the *A. thaliana* dataset contained 3,421,779,258 sequenced bases in 300,071 reads (~25.22x coverage of the 135.67Mb reference sequence), and the *D. melanogaster* dataset contained 4,617,842,308 sequenced bases in 663,784 reads (~32.39x coverage of the 142.57Mb reference sequence). We obtained the reference genome sequence for *E. coli* strain K12_MG1655 from NCBI; all other reference genome sequences were obtained from Ensembl (Release 95). Accession numbers are provided under ‘Resource Availability.’

We simulated 150bp paired-end Illumina DNA libraries with the software ART [20] and ONT DNA libraries with the software NanoSim (version 2.0.0; [21]). Both software programs utilize an assembled sequence to generate simulated libraries with read profiles similar to an empirically generated DNA library. NanoSim additionally requires real ONT flowcell data to simulate unique, organism-specific mismatch, insertion and deletion rates. ART libraries were simulated to a coverage of 300x with a fragment standard deviation of 50. NanoSim libraries mirror the characteristics of the real ONT flowcells including read length distributions. We used NanoSim to simulate 500,000 *E. coli* reads (943x coverage; read N50 8,158), 2,000,000 *C. elegans* reads (336x coverage; N50 21,559), 5,000,000 *A. thaliana* reads (420x coverage; N50 19,577) and 10,000,000 *D. melanogaster* reads (219x coverage; N50 11,955). The sequence read N50 indicates 50% of the total sequenced nucleotides are in reads that long or longer.

### C. remanei *sequence libraries*

Output for each flow cell used for each strain is summarized in STable 1. Based on an estimated genome size of 125Mb [22] we generated 76x-156x coverage of each strain with sequence read N50 8,203-22,891bp.

## Genome assembly

### Assembly of Simulated Data

Genome sequences were assembled using three approaches. We first assembled the simulated ONT read sets with Canu v1.9 [18]. Each genome was assembled at decreasing coverage depths until the assembler failed to produce an assembly with the given data. In order to minimize the influence of individual reads and stochastic assembly artifacts, each read set was generated by selecting a random subset of the full simulated dataset. The second approach used the same subsets of ONT data, but with Flye 2.8 [23] as the assembly software.

The long-read datasets that performed the best were paired with 2×150 simulated paired-end Illumina data and assembled using MaSuRCA version 3.3.9 [24]; coverage depths were adjusted for both datasets to better understand the effects of increasing or decreasing coverage on the final assembly. Here, we retained the ONT dataset to maximize our ability to draw parallels between assembly approaches. For example, the minimum *C. elegans* ONT dataset that assembled with Canu [18] was an average of ~60x coverage across the genome and this read set was used in the MaSuRCA trials with 50x and 100x Illumina coverage, respectively.

We also used Flye [23] to assemble read sets that had been corrected using the ‘canu – correct’ function. Canu correction uses all-versus-all overlap information to correct each individual read [18]. In addition to the consensus information, Canu employs two filters to avoid biases due to sequence quality or repeats. From read-length estimates, Canu uses the longest available reads up to the user-specified coverage depth for correction [18]. By default, this is set to 40x coverage. Assembling these datasets with both Canu and Flye allowed us to compare the effectiveness of the error correction modules within each software package. We also assembled libraries with reads selected by the Canu correction module but not subject to correction to test the influence of error correction vs. read length on assembly.

The most contiguous assemblies from the long-read only and hybrid categories were error corrected using Pilon version 1.23 [25] to determine the effect of short-read polishing on the accuracy of the draft assemblies. Each simulated assembled sequence was polished with the entire simulated paired-end data set. Four rounds of polishing were completed for each assembly with statistics measured after each round with QUAST v5.0.2 [26] and BUSCO v4.0.3 [27].

### Assembly of ONT Nematode Data

Preliminary assemblies were created for each strain by first correcting the complete library with canu –correct (Canu v2.0) [18]. Corrected reads from Canu were then assembled using Flye 2.8 [23]. The draft assembly for each strain was polished using Pilon v1.23 [28] and Illumina paired-end reads. All data sets contained a small sub-set of contaminants from the nematode microbiome that were removed from the final assemblies [29, 30].

## Analyses

### Assessing the quality of assembled simulated sequences

We measured genome statistics relative to the reference sequence of each organism with QUAST [26]. We assessed contiguity and accuracy of the assembled sequence through eight statistics:

(1) **NG50** is a size median statistic and indicates that 50% of the expected assembled genome sequence (where the expectation is based on a known reference) is contained in contiguous sequences that large or larger.
(2) **NGA50** is a similar size median but indicates that 50% of the expected assembled genome sequence that aligns to the reference genome is contained in contiguous sequences that large or larger.
(3) **LG50** is the number of linkage groups or contiguous assembled sequences containing 50% of the expected assembled genome sequence.
(4) **LGA50** is similar but again measured in the portion of the assembled sequence that aligns to the reference genome.
(5) **Genome fraction (%)** is the fraction of reference genome captured in the assembled sequence.
(6) **Duplication** measures the fraction of reference genome found multiple times in the assembled sequence.
(7) **Mismatches** is the number of mismatched nucleotides per 100,000 nucleotides or base pairs (bp).
(8) **Indels** are the number of insertions and/or deletions per 100,000 nucleotides.

For our metazoan organisms we also used the software package BUSCO (version 4.0.3; released February 2020) to search for a set of unique genes expected to be conserved in single copy in an evolutionarily related group of organisms [27]. Below, we describe the analyses of each organismal dataset.

### Escherichia coli

The *E. coli* genome is relatively small at 4.64Mbp and contains few repeats (87.8% of the genome codes for proteins; [31]). Assembly of 50,000 ONT sequences (~62x coverage; N50 8,350bp) with Canu [18] and Flye [23] resulted in circular contigs with high contiguity and few mismatches, insertions, and deletions (Fig. 2A-D). At low ONT coverage (~19x) the assemblies produced by Canu and Flye were contained in 6 and 9 contigs, respectively. Both assembled sequences had high rates of mismatches, insertions and deletions when compared with sequences assembled from high coverage libraries (Fig. 2C-D). Increasing ONT coverage beyond 62x resulted in poor contiguity for Canu assembled sequences (Fig. 2A) and increased mismatches for Flye assembled sequences (Fig. 2C). Assembly of hybrid Illumina-ONT libraries with MaSuRCA [24] resulted in sequences with variable contiguity and accuracy. Two of the tested hybrid sets were able to assemble the genome into a single contig but the library with the highest Illumina coverage (300x) and highest ONT coverage (227x) assembled into two contigs. MaSuRCA was unable to perform as well as either Canu or Flye when given the same long-read dataset. For instance, the top performing Canu and Flye runs used 50,000 ONT reads (~62x coverage); the same reads passed through MaSuRCA produced 2 contigs, regardless of Illumina coverage (Fig. 2B).

**Figure 2.**
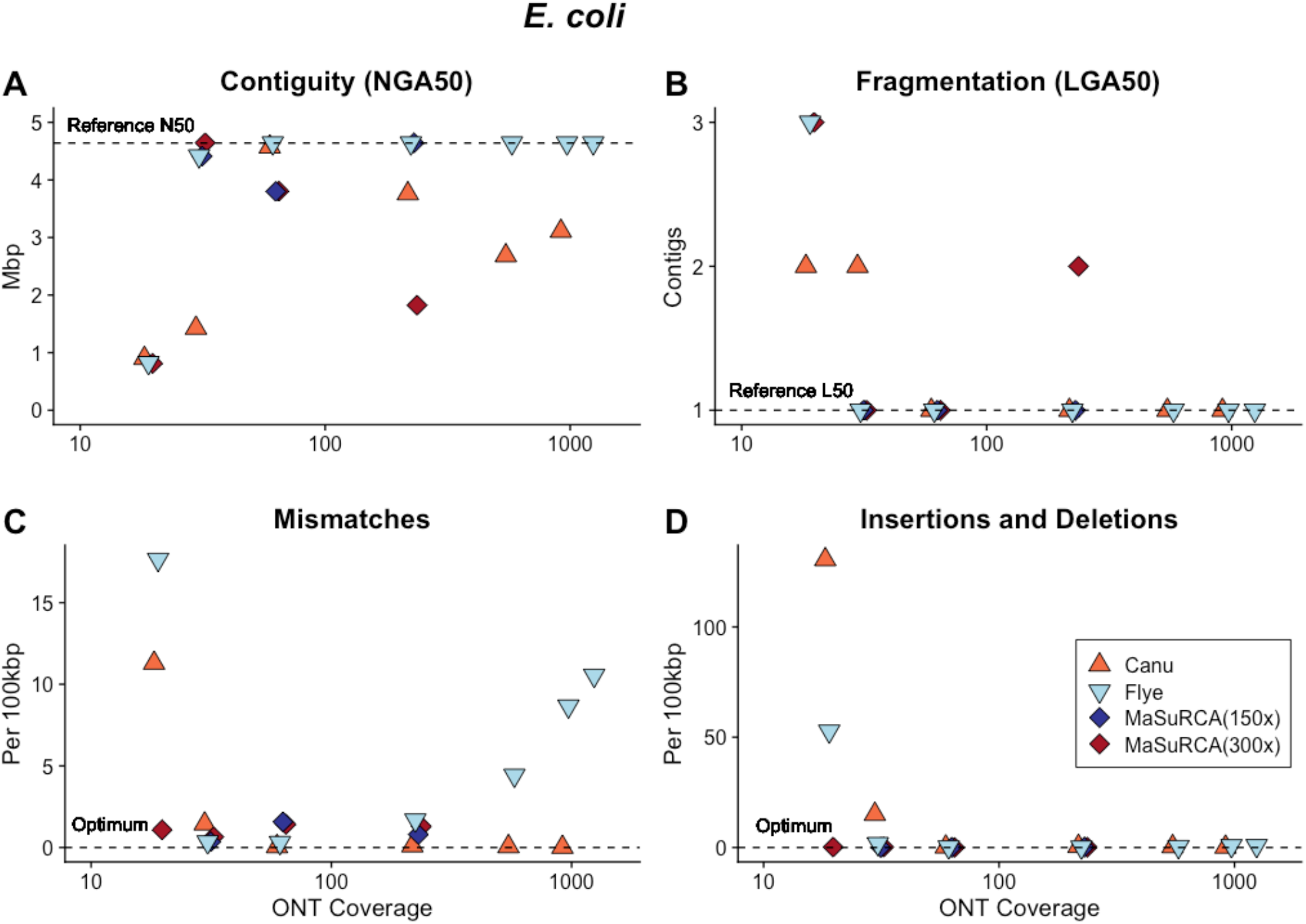
The *E. coli* genome is relatively small at 4.64Mbp and less complex when compared with metazoan genome sequences. Assembly of ONT libraries at relatively high coverage (>60x average sequence depth) with Canu and Flye results in assembled sequences with (A) high contiguity; (B) low contig number converging on a single chromosome; (C) few mismatches; and (D) few insertion/deletion errors (indels) when compared with the reference sequence. The influence of sequencing coverage on contiguity or fragmentation was not monotonic for Canu or MaSuRCA assembled sequences. The relationship between sequencing coverage and accuracy was not monotonic for Flye assembled sequences.

After polishing the assemblies with Pilon [25], the Canu-assembled sequence had fewer mismatches and indels per 100 Kbp than the Flye assembled sequence or an assembly produced by Flye using reads that had first been corrected with Canu (SFig. 1A-C). Despite its dependence on higher accuracy Illumina sequences the MaSuRCA assembled sequence had higher numbers of mismatches, insertions and deletions both pre- and post-polishing (SFig 1B). We used MaSuRCA to assemble a set of paired end Illumina sequences to test the influence of adding ONT data to the assembly process. The resulting sequence was 98.76% of the *E. coli* genome and was fragmented into 74 contiguous pieces. The accuracy, 1.7 mismatches and 0.04 indels per 100 Kbp, was improved when compared with the hybrid assembled sequences. This indicates that the inclusion of ONT data can introduce errors in hybrid approaches that cannot be corrected with post-assembly polishing.

### Caenorhabditis elegans

The *C. elegans* genome is 100.8 Mbp [32] contained in a single X chromosome and five autosomes [33]. *C. elegans* is a diploid self-fertile hermaphrodite with low levels of genetic diversity [34] and approximately 16% of the genome is repetitive [35]. Canu, Flye, and MaSuRCA assembled sequences plateaued in contiguity and accuracy above ~70x coverage (Fig. 3A-D). The read N50 for these datasets ranged from 20,742 to 22,840. Both Canu and Flye assembled chromosome-scale contigs with low rates of mismatches, insertions and deletions. The MaSuRCA [24] hybrid assembly approach did not perform as well, even with high ONT coverage (Fig. 3A,B).

**Figure 3.**
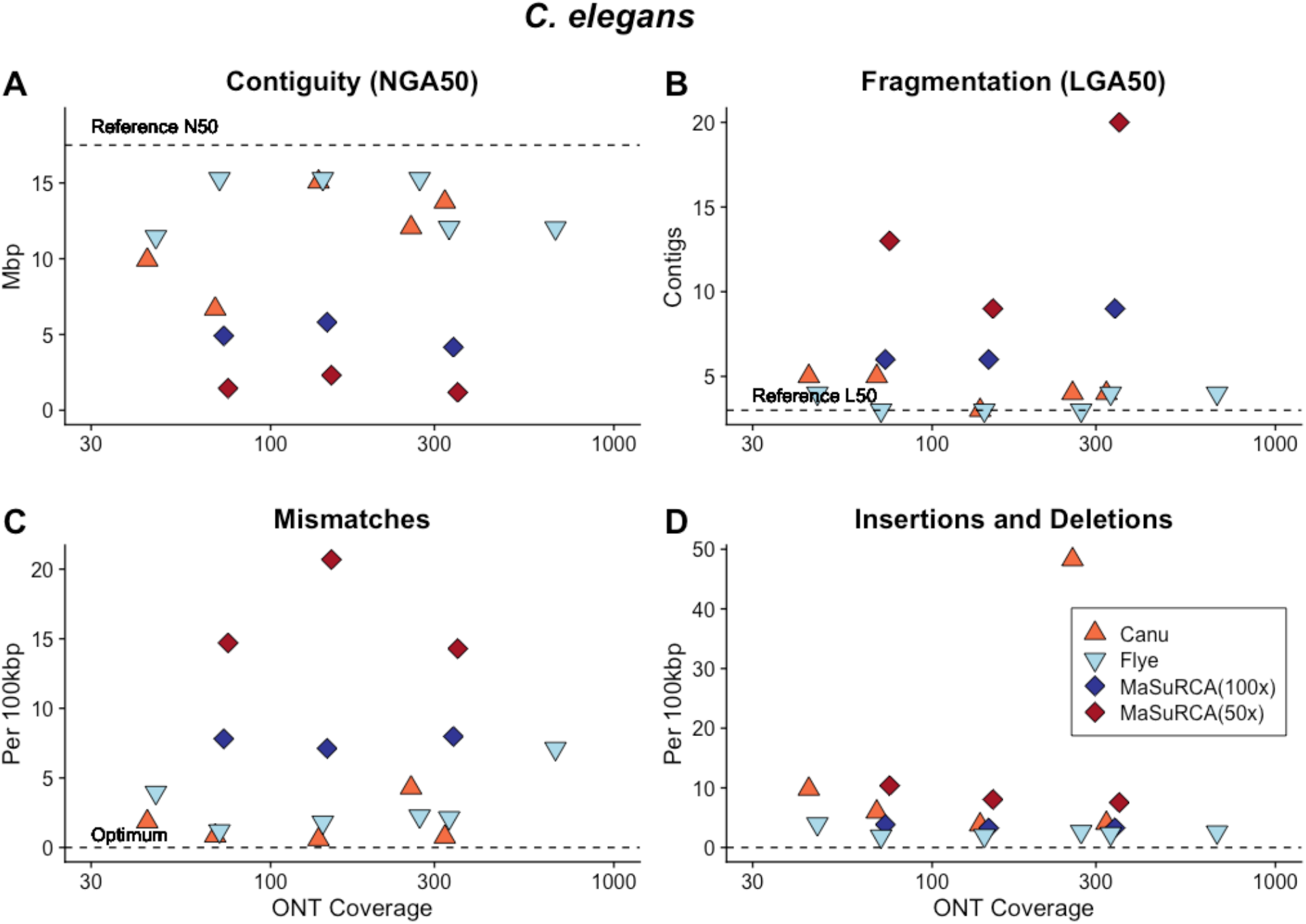
Both Canu and Flye assembled libraries with >70x ONT coverage into genome sequences with A) high contiguity; B) low fragmentation; C) few mismatches; and D) few insertions and deletions when compared with the *C. elegans* reference sequence. In comparison MaSuRCA performed poorly across a range of ONT sequencing coverages.

To improve these assemblies we experimented with polishing the Canu and Flye assembled sequences, Additionally, we used Flye to assemble the Canu corrected dataset (“Corrected” in Fig. 4) and used Flye to assemble the reads selected by the Canu correction module but not corrected (“Selected” in Fig. 4). The dataset corrected with Canu and assembled with Flye had a read N50 of 28,641bp and produced 6 chromosome-scale contigs with low numbers of mismatches, insertions and deletions that further decreased after polishing (Fig. 4A-D). Flye assembly of the same dataset prior to correction resulted in an assembly with higher fragmentation (Fig. 4B) and lower accuracy (Fig. 4C-D). This dataset had a read N50 of 29,404bp, indicating that both read length and correction contribute to assembly quality. The assembly graph for the Canu corrected, Flye assembled dataset (Fig. 5) shows 6 chromosome-scale contigs with smaller disjunct assembled sequences [36].

**Figure 4.**
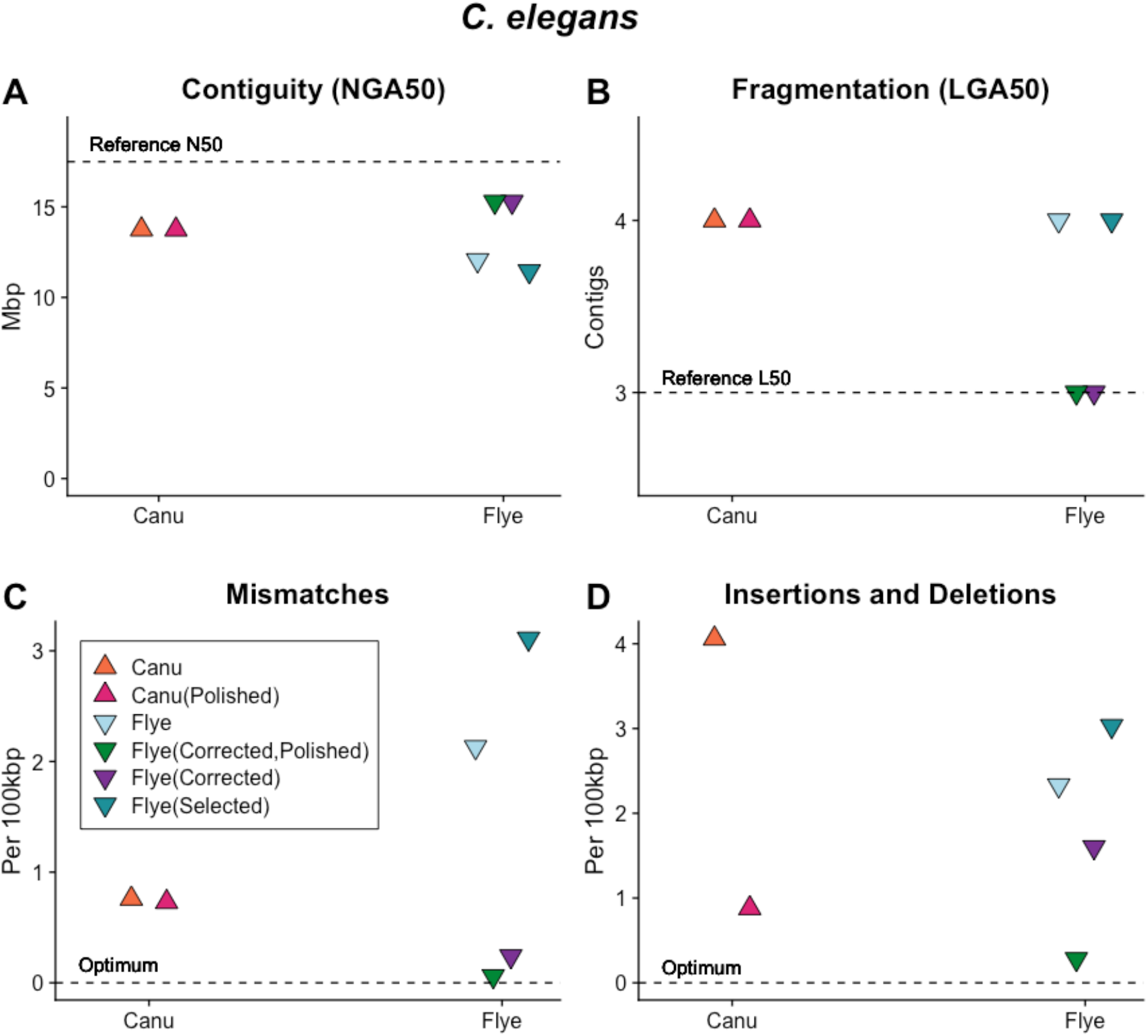
The relative performance of Canu and Flye assembly software with raw datasets, datasets with reads selected for length (‘Selected’), reads selected for length and pre-corrected (‘Corrected’ in the figure legend) and sequences post-assembly polished with Illumina short reads (‘Polished’ in the figure legend) as measured by A) genome contiguity; B) mismatches and C) insertions and deletions relative to the *C. elegans* reference sequence. Canu assembly of *C. elegans* ONT libraries produces assembled sequences with high contiguity, low fragmentation, few mismatches and relatively more insertions and deletions that are eliminated with post-assembly polishing. The performance of Flye was increased through read selection, correction and post-assembly polishing with the highest-quality *C. elegans* sequence produced through Canu correction, Flye assembly and post-assembly polishing. The dataset with reads selected for length did not assemble as well as the dataset with reads selected and corrected, indicating that correction is an important step in assembly.

**Figure 5.**
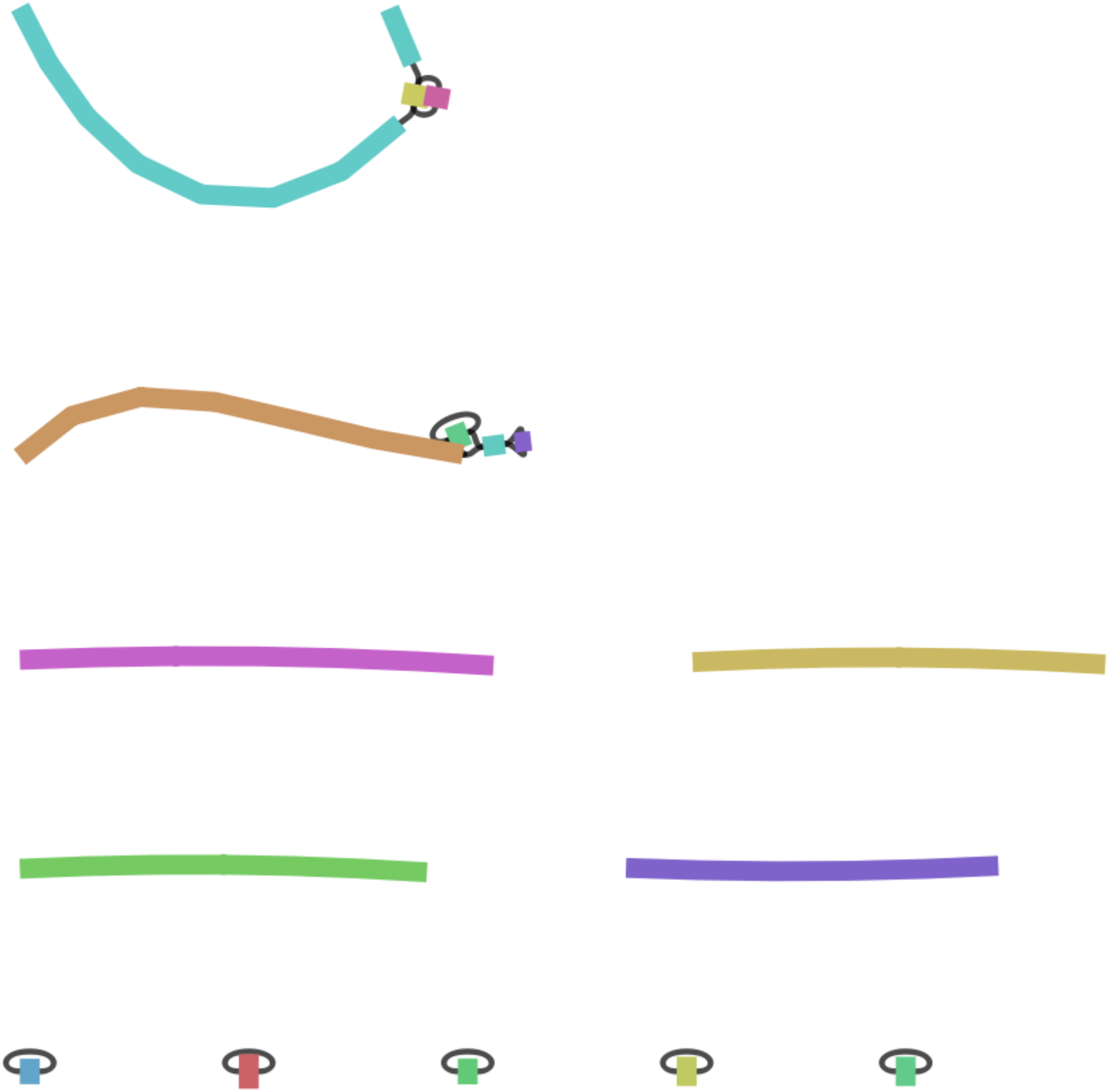
The assembly graph for the Flye assembled Canu selected and corrected *C. elegans* dataset shows 6 large sequences corresponding to the 6 chromosomes of *C. elegans* with 5 small unconnected sequences. Two of the large sequences contain unresolved ‘bubbles’ in the assembly graph.

We used QUAST [26] to identify annotated features in our assembled sequences to compare each software’s ability to assemble different classes of genomic elements. We used the WS277 annotations provided by WormBase Parasite and identified exons, introns, pseudogenes, repeats, small RNAs, transposable elements (TEs) and untranslated regions (UTRs). The worst performing assembly had >99.67% representation of exons, introns, small RNAs and UTRs and >98.85% representation of repetitive sequences like pseudogenes, repeats and TEs (Table 1). The Flye assembled sequences had the best representation of exons, introns, small RNAs and UTRs while the Canu assembled sequences had the best representation of repetitive pseudogenes, repeats and TEs.

**Table 1.** The proportion of genomic features that QUAST identified in the assembled genome sequence. The best performing software and dataset combination is highlighted in bold for each genomic feature. Flye reliably assembled exons, introns, small RNAs and UTRs while Canu performed best when assembling pseudogenes, repeats and transposable elements. Polishing the assembled genome sequence with Illumina short reads did not affect the inference of genomic features.

| Assembly Software | Dataset | Exons | Introns | Pseudo-genes | Repeats | Small RNA | TEs | UTRs |
| --- | --- | --- | --- | --- | --- | --- | --- | --- |
| Flye | Selected | **0.9988** | **0.9993** | 0.9954 | 0.9967 | **0.9985** | 0.9941 | 0.9996 |
| Flye | Corrected | **0.9988** | **0.9993** | 0.9954 | 0.9971 | 0.9984 | 0.9942 | **0.9997** |
| Flye | Corrected, Polished | **0.9988** | **0.9993** | 0.9954 | 0.9971 | 0.9984 | 0.9942 | **0.9997** |
| Canu | Corrected, Polished | 0.9986 | 0.9986 | **0.9986** | **0.9981** | 0.9981 | **0.9978** | 0.9988 |
| MaSuRCA |  | 0.9967 | 0.9973 | 0.9885 | 0.9897 | 0.9972 | 0.9887 | 0.9975 |

### Drosophila melanogaster

The 180Mb genome of the diploid fruitfly *D. melanogaster* presents multiple challenges for sequencing and assembly including approximately 60Mb of repetitive heterochromatin [37]. The genome sequence is contained in a large X chromosome, a small (“dot”) Y chromosome and 3 autosomes. Assemblies of the simulated ONT *D. melanogaster* libraries repeated patterns seen with the *C. elegans* dataset; the most contiguous Canu assembly was produced with ~113x ONT coverage and a read N50 of 11,679 (~16Gbp of data; Fig. 6A,B). This assembly produced 145 contiguous pieces but many of these were small and the LGA50 was 4 contigs (Fig. 6B). The Flye assembly of the same dataset was much less contiguous, producing 482 contigs. When the dataset was corrected with Canu then assembled with Flye the assembled sequences had high accuracy (SFig. 2).

**Figure 6.**
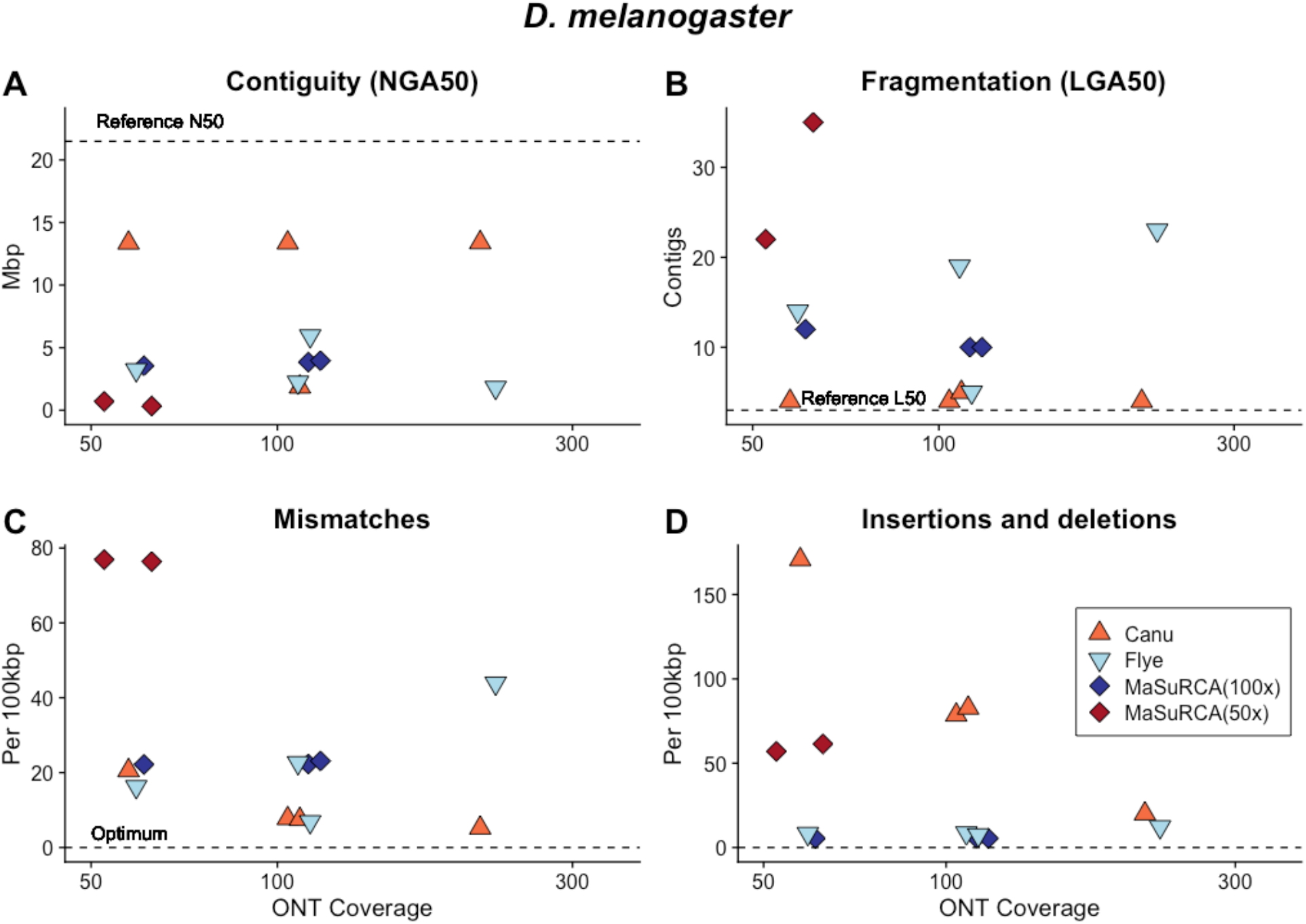
The *D. melanogaster* genome presents multiple challenges for genome sequencing and assembly including approximately 60Mb of repetitive heterochromatin. Canu assembly of ONT libraries with >60x sequencing depth produced a sequence with (A) high contiguity; (B) low contig number; (C) few mismatches and (D) few insertions and deletions (indels) when compared with the reference. The accuracy of the Canu assembled sequence increased with ONT library coverage. Flye assembly of the same ONT libraries produced sequences with relatively less contiguity, more fragmentation, more mismatches and fewer insertions and deletions. MaSuRCA assembly of the same ONT sequences resulted in fragmented sequences with similar error levels with compared with the reference sequence.

The top performing Canu assembly contained 91% of the metazoan genes expected to be conserved in single copy with BUSCO [27] prior to polishing and 94% after four rounds of Pilon with paired-end reads (SFig. 3). In comparison, the *D. melanogaster* reference sequence contains 94.1% of the expected single copy metazoan genes. Additional data decreased contiguity with a slight increase in accuracy and NGA50 (Fig. 6 A,D). Hybrid MaSuRCA [24] assemblies for *D. melanogaster* performed markedly worse than Canu assemblies of ONT data (Fig. 6B). The top performing MaSuRCA assembly produced 220 contigs with 93.9% of the expected metazoan genes identified [27].

### Arabidopsis thaliana

The 135Mbp genome sequence of the self-fertile plant *A. thaliana* is contained in 5 autosomes [38]. *A. thaliana* has undergone at least two rounds of whole genome duplication [39] and contains large tracts of highly similar genomic regions. Assembly of the ONT libraries with Canu [18] and Flye [23] resulted in sequences with lower contiguity and higher fragmentation when compared with the other model organisms, even when the library contained ~420x coverage and a read N50 of 19,577bp (56.7Gbp of sequenced nucleotides). Canu produced 30 contiguous pieces (LGA50 5 chromosomes) with 98.8% of the expected Viridiplantae genes (Supplementary Data) identified prior to polishing [27] and Flye produced 26 contigs and 98.8% of the expected Viridiplantae genes. The Flye assembly contained many more mismatches and indels than its Canu counterpart, but this discrepancy was alleviated with Pilon polishing (SFig. 5). Following polishing with Pilon [25], the Flye assembled sequences contained 99.1% of the expected Viridiplantae genes [27], matching that of the TAIR10 reference genome for *A. thaliana*. Interestingly, our combined approach of long-read methods did not work with *A. thaliana* datasets and neither Flye nor Canu was able to assemble a draft genome with <40x coverage.

The hybrid MaSuRCA assemblies for *A. thaliana* performed the best of the three eukaryotes. The top performing MaSuRCA [24] assembly produced 45 contiguous pieces with 98.8% of the expected Viridiplantae genes identified in the sequence [27]. This was also brought up to 99.1% after polishing with Pilon (Supplementary Data).

### *Caenorhabditis remanei* and *Caenorhabditis latens*

We used our simulated data to create a protocol for *de novo* sequencing and assembly for 3 strains of the nematode *C. remanei* [40] and one of the closely related nematode *C. latens* [41]. Both *C. remanei* and *C. latens* are obligate outcrossing species with high levels of nucleotide diversity [42] that have hobbled previous assembly attempts [35, 43]. The worm’s small size means that pools of individuals harboring both individual and population-level diversity must be sacrificed for DNA sequencing and assembly. We prepared high-molecular-weight DNA extracts and generated ONT libraries for an outbred *C. remanei* strain EM464, two inbred laboratory strains PX356 and PX439 and the inbred laboratory *C. latens* strain PX534. We tested library preparation and computational protocols with the inbred *C. remanei* strain PX356 and used these to generate assembled sequences for the other *C. remanei* and *C. latens* strains.

The most contiguous *C. remanei* PX356 assembly was achieved by using Canu [18] to correct the entire ~156x coverage ONT dataset prior to assembly with Flye [23]. After correction, the ~40x coverage dataset had a read N50 of 21,340bp and resulted in an assembled sequence contained in 163 contigs with 89.1% of the expected conserved nematode genes [27]. Decontamination with Blobtools2 [29] and SIDR [30] resulted in a final assembly for PX356 with 124,499,812bp contained in 70 contigs with an N50 of 6.2Mbp and an N90 of 1.93Mbp contained in just 18 large contigs, approximately 3 per chromosome. We compared our results (Fig. 7A-D) with a recently published sequence for the related *C. remanei* strain PX506 [22] generated with Pacific Biosystems sequencing and chromatin conformation capture to produce chromosome-scale sequences [44].

**Figure 7.**
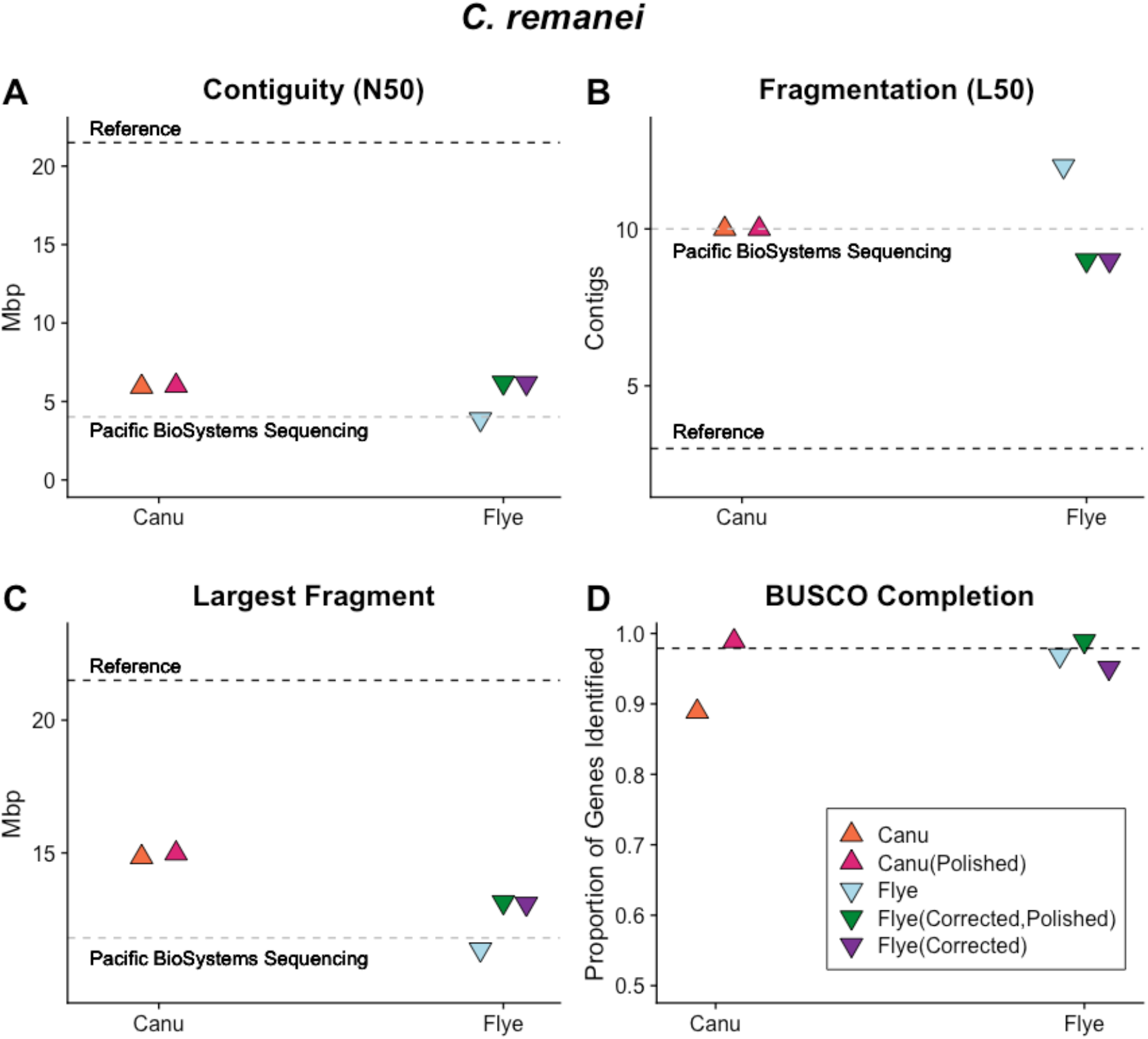
Both Canu and Flye produced assembled sequences with A) contiguity; B) fragmentation; C) the largest fragment and D) BUSCO gene completeness comparable to Pacific BioSystems sequencing after read correction and post-assembly polishing.

Canu-only and Flye-only assemblies contained more contigs and a smaller N50 than the combined approach described above (Fig. 7A-D). The assembly graph (Fig. 8) shows several large contiguous fragments but also multiple regions difficult to disentangle and smaller disjunct fragments. Analyses with GenomeScope [45] suggested ~0.7% of the genome remains heterozygous (SFigure 6). Assemblies with subsamples of the long-read data were increasingly fragmented with decreasing coverage. Flye assemblies with initial coverages of 102x, 70x, and 52x produced 183, 242, and 267 contigs respectively. A MaSuRCA-hybrid approach [24], using 102x ONT coverage and 450x paired-end coverage yielded 336 contigs and 96.6% of BUSCO single-copy genes.

**Figure 8.**
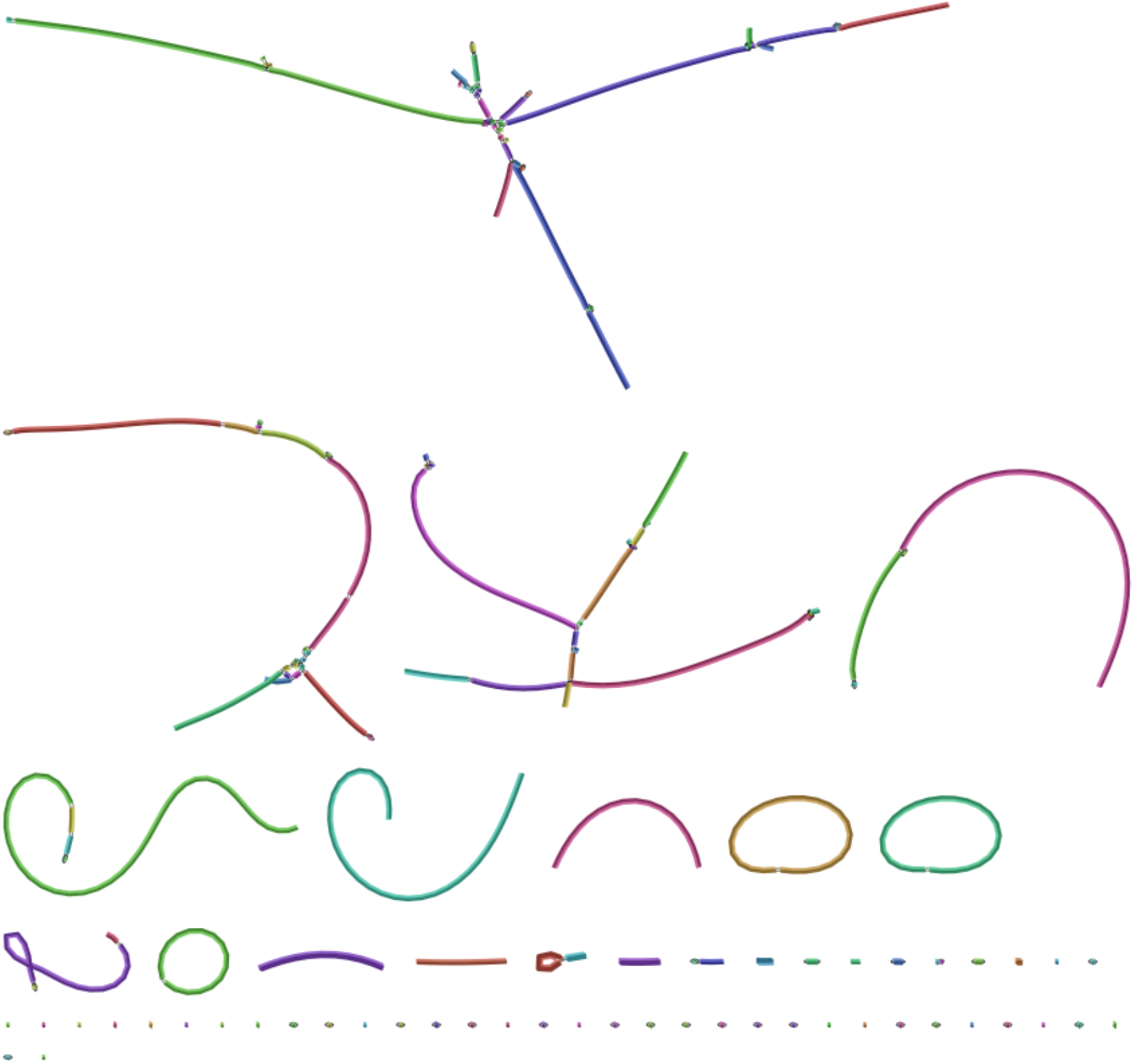
The *C. remanei* PX356 assembly graph shows multiple unresolved regions, bacterial contaminants (visible as circular chromosomes) and small fragments.

BUSCO identified 97.7% (increased from 89.1%) of the expected conserved single copy genes following 4 rounds of polishing using Pilon [28] with Illumina paired end reads at ~225x average depth (Fig. 6). We tested the influence of Illumina coverage on polishing and found that 97.6% of the expected conserved nematode genes could be identified after polishing with just ~20x Illumina coverage (STable 3), indicating that a large amount of data is not necessary to correct the majority of errors in an assembly. However, this was achieved after three successive rounds of error correction with the Pilon software package [28] utilizing the same Illumina DNA sequence reads. This assembled sequence is less fragmented and contains a higher percentage of conserved nematode genes (Fig. 6) when compared with the previously published assembly for *C. remanei* PX356 produced with Illumina libraries and a genetic linkage map [35].

We generated assembled genome sequences for *C. remanei* PX439, *C. remanei* EM464, and *C. latens* PX534 using the ‘best practices’ protocols developed through simulations and our *C. remanei* PX356 experimentation. Our *C. remanei* PX439 dataset had an ONT coverage depth of ~120x with a read N50 of 18,233bp. After Canu correction the read N50 increased to 38,434bp and we assembled a 132,046,643bp genome sequence contained in 53 contigs (N50 12.6Mbp). The N90 for this assembled sequence was 2.27Mbp contained in 12 large contigs, approximately 2 per chromosome.

*C. remanei* EM464 is outbred and retains high levels of heterozygosity. We did not expect to assemble a cohesive genome sequence but generated ONT libraries and an assembled sequence to identify the distribution of allelic variants in the population. The ONT library had a coverage depth of ~76x and a read N50 of 22,891bp. The Canu corrected dataset used for Flye assembly had an N50 of 34,951bp. Flye produced a 138,626,018bp genome sequence contained in 320 contigs (N50 3.4Mbp). The flow-cytometry estimated genome size of *C. remanei* is 124-131Mbp and we estimate that 5-12% of the assembled sequences are allelic variants. The *C. latens* PX534 dataset had a coverage depth of ~119x and a read N50 of 14,759bp. After Canu correction the read N50 of the dataset 34,591bp and we assembled a 120,367,464bp genome sequence in 97 contigs with an N50 of 13.6Mbp.

## Discussion

We have found that Canu [18] and Flye software packages [23] produce high-quality contiguous draft assemblies from ONT libraries for a broad range of organisms. In many cases, Flye slightly outperforms Canu while being much faster and using fewer computational resources [13, 46], however there are many instances in our results where the best assembly is produced by first error correcting the data with the Canu correction modules, then assembling the corrected reads with Flye. Given the same set of corrected reads, the Flye assembler regularly creates a better assembly than Canu. This finding highlights the need for performing multiple assembly strategies to identify the top performing strategy for each dataset.

We found the hybrid assemblies produced by MaSuRCA [24] contained a higher proportion of expected conserved genes when compared with the unpolished long-read only assemblies and had fewer mismatches and insertion/deletion errors. However, given the same amount of ONT data, both Canu [18] and Flye [23] assembled more contiguous genome sequences. The long-read only assemblies also produced larger NG50 values and contained a larger fraction of the expected genome when compared to hybrid MaSuRCA assemblies. The shortfalls of long-read only assembly can be overcome by ‘polishing’ with low coverage (~20x) Illumina DNA sequences using a software such as Pilon [28] to error correct the draft assemblies. We found that polishing played a larger role in increasing accuracy of the real-world data compared to the results seen with simulated datasets. These methods improved the accuracy of Canu or Flye assemblies to be on par with, or better than, those produced by MaSuRCA.

These highly contiguous assemblies were achieved with relatively high sequencing depth, at least 100x coverage across the genome. Although ONT can theoretically produce megabase-sized reads, in reality, many of the sequence reads in ‘real’ projects are shorter due to fragmentation that occurs during DNA purification and library preparation. ONT libraries may have many more small reads and >100x ‘real’ coverage may be necessary to achieve >40x coverage with reads >25 Kbp and highly contiguous assemblies. Increasing read N50 in a library through improvement of DNA isolation and library preparation methods reduces the necessary coverage to produce a high-quality assembly [47]. Therefore, researchers should adjust their coverage goals based on the quality of libraries they are able to reliably produce.

Our findings with simulated datasets were supported by all real-world datasets tested. *Caenorhabditis remanei* PX356, PX439 and *C. latens* PX534 all have previously published genome assemblies generated with paired-end Illumina sequences [35]. Our assemblies proved to be at least one order of magnitude more contiguous than any of the previous assemblies, highlighting the ability of ONT long reads to drastically improve contiguity. Outbreeding Caenorhabditis are resistant to inbreeding [48] and previous assembly attempts resulted in over-assembly due to residual heterozygosity [49] or under-assembly. Despite this, we were able to finalize a highly contiguous and complete assembly for these nematodes. The strains we targeted are closely related and interfertile [50] yet their assembled genome sequences vary up to 10% in length. This level of structural variation between closely related species highlights the need for *de novo* assembled sequences in molecular evolution research.

Our *de novo* assembled *Caenorhabditis* sequences are contiguous and achieved BUSCO scores greater than those of Illumina-only assembled sequences (STable 4). The findings we present here clearly demonstrate existing genome sequences can be improved by adding in ONT datasets. As sequencing and assembly technologies continue to improve and drop in cost, many researchers will find published datasets can be updated to further improve downstream analyses.

## Potential Implications

In light of our findings, we suggest that long-read data be prioritized when undertaking *de novo* genome assembly projects. Our results indicate that an assembly with sufficient ONT long read coverage will be highly contiguous and polishing with Illumina data can achieve high levels of accuracy. For situations where high-coverage ONT libraries are not feasible, MaSuRCA-assembled [24] Illumina and ONT read sets can produce reliable draft sequences. However, the quality and contiguity of the assembled sequence is determined by Illumina read depth and effort should be made to increase Illumina read depth, even if it is at the expense of ONT sequences. MaSuRCA-assembled Illumina sequences have fewer mismatches and insertion/deletion errors when compared with MaSuRCA-assembled ONT and Illumina hybrid read sets, indicating that the inclusion of ONT sequences introduces errors. We suggest that error correction with Illumina DNA sequences and the Pilon software package [25] is a necessary finishing step in any assembly project utilizing ONT data.

Our results demonstrate that near-chromosome-level genome sequences are achievable with sufficient ONT data. However, chromosome-level genome assemblies are often not necessary to address many research questions, particularly those focused on small numbers of genes or phylogenomic information. Researchers should approach genome sequencing by first determining what genome-completion level will be sufficient for their research goals. To aid in this approach, we hope our study will help researchers determine the amount of sequencing effort, and the sequencing approaches, that will best suit their needs.

## Resource Availability

### Lead Contact

Janna L. Fierst

### Materials Availability

This study did not generate new unique reagents.

### Data and Code Availability

*E. coli* genome sequence GCF_000005845.2; ONT read set SRR8154670

*C. elegans* genome sequence GCA_000002985.3; ONT read set ERR2092776

*A. thaliana* genome sequence GCA_000001735.1; ONT read set ERR2173373

*D. melanogaster* genome sequence GCA_000001215.4; ONT read set SRR6702603

*C. remanei* PX356 BioProject PRJNA248909

*C. remanei* PX439 BioProject PRJNA248911

*C. remanei* EM464 PRJNA562722

*C. latens* PX534 BioProject PRJNA248912

Full datasets are available at https://doi.org/10.5061/dryad.3r2280gd2

Bioinformatic scripts and workflows are available at https://github.com/jmsutton2

## Methods

### DNA Extraction and Sequencing

Nematodes used for genomic sequencing were grown on two 100mm NGM plates [51] seeded with *E. coli* OP50. Worms were harvested by washing plates with M9 minimal media into 15mL conical tubes. Samples were rocked on a tabletop rocker for 1 hour before being centrifuged to pellet worms. The supernatant was removed and tubes refilled with sterile M9, mixed and pelleted by centrifugation again. This process was repeated five times or until the supernatant was clear after centrifugation. The pellet was moved to 2mL tubes and frozen at – 20°C until extraction. Worm pellets were allowed to thaw to room-temperature then flash frozen in liquid nitrogen and repeated three times. Worm pellets were placed in 1.2mL of lysis buffer solution (100mM EDTA, 50mM Tris, and 1%SDS) and 20μL of Proteinase K (100mg/mL). Tubes were then placed on a 56°C heat block for 30 minutes with shaking. Genomic DNA was then isolated using a modified phenol-chloroform extraction[52]. DNA concentrations and purity were measured with a Qubit 4 (Life Technologies, Carlsbad, CA USA) and NanoDrop^®^ 1000 spectrophotometer, respectively (Thermo Fischer Scientific, Waltham, MA USA). Extracts were visualized on a 0.8% agarose gel to verify high-molecular weight gDNA. DNA size selection was carried out using the Short Read Eliminator Kit from Circulomics Inc. according to manufacturer guidelines (Baltimore, MD USA).

DNA libraries were prepared for each sample using the SQK-LSK109 ligation sequencing kit and loaded on to R9.4.1 RevD flow cells. The recommended protocol from ONT was modified by replacing the first AmpureXP bead clean step with an additional treatment with the Short Read Eliminator Kit. Approximately 700ng of gDNA from each library was loaded on to a flow cell sequenced for 48 hours on a GridION X5 platform with basecalling performed by Guppy v.4.0.11. ONT reads for each strain were trimmed of adapters using Porechop (https://github.com/rrwick/Porechop) with the –discard_middle flag to remove chimeric reads.

### Assembly and Decontamination of Real Datasets

We evaluated contaminants in the *C. remanei* PX356, *C. remanei* PX439, *C. remanei* EM464 and *C. latens* PX534 draft assemblies with Blobtools2 [29] and SIDR [30]. Blobtools2 uses phylum-level taxonomic assignment, read coverage depth and GC content to identify non-target contigs within the assembly. To prepare data for visualization with BlobToolKit, the ONT reads were aligned to the initial assemblies using minimap2 v2.17 [53] and sorted using SAMtools v1.8 [54]. A reference database for taxonomic identification of scaffolds was created using blastn 2.2.31 [55] using the nt database from the National Center for Biotechnology Information. An additional database was generated using diamond tblastx [56]. Contigs that were >10,000bp and taxonomically identified as *Nematoda* were retained in the assembly.

SIDR [30] utilizes ensemble based machine learning to train a model capable of discriminating target and contaminant contigs based on measured predictor variables. To prepare data for SIDR decontamination Illumina DNA and RNA sequence reads were aligned to the draft assembly with bwa [57] using the BWA-MEM algorithm. We used bbtools/bbmap software [58] to calculate the average sequence coverage, length, GC, bases covered in alignment, RNA average sequence coverage and bases covered in RNA alignment for each contig. We used these data to train a bootstrap aggregated (bagged) decision tree model based on the blast identification of contig origin and used this model to predict the origin, target or contaminant of each contig. SIDR allowed us to assign probable *Nematoda* origin to contigs that lacked blast identification. We discarded contaminants common to *Nematoda* genome assemblies including E. coli, Pseudomonas, Serratia and Stenotrophomonas [59]. We also identified one large 4.57Mb *E. coli* contig in the *C. latens* PX534 that had been labeled *Nematoda* by BlobToolKit and removed it from the *C. latens* PX534 assembled sequences.

### Heterozygosity analyses

We assessed heterozygosity using the software Jellyfish version 2.2.4 [Marcais 2011] and GenomeScope [Ranallo-Benavidex 2020]. Jellyfish is a fast, memory efficient kmer counting tool that searches for occurrences of unique kmers in raw sequence reads based on user selected size. GenomeScope estimates genome heterozygosity, genome size, and repeat content using a k-mer based statistical approach. Jellyfish was run using the default settings and a kmer size of 21. This kmer size was selected based on recommendation from the maker of Jellyfish and should be sufficient to obtain all unique kmers present. A histogram file of all kmer occurrences was created by Jellyfish and then uploaded into the online interface of GenomeScope. GenomeScope was run using default settings with a kmer length of 21 and a read length consistent with each sample’s sequencing parameters.

### Evaluation

We used the software BUSCO version 4.0.1 [27] to identify conserved gene sets. Briefly, BUSCO searches assembled DNA sequences for a set of unique genes that are expected to be conserved in single copy in an evolutionarily related group of organisms. We used BUSCO version 4.0.1 [27] and the datasets Nematoda_odb10 for *C. elegans*, *C. remanei,* and *C. latens*, Metazoan_odb10 for *D. melanogaster*, and Viridiplantae_odb10 for *A. thaliana*. We also measured BUSCO completeness with the Diptera_odb10 for the *D. melanogaster* assemblies but found that in some instances our assembled sequence contained a greater proportion of conserved genes than the reference sequence. We chose to focus on the Metazoan_odb10 for *D. melanogaster* and present both sets of statistics in the Supplementary Data. BUSCO genes in each assembly were classified as single copy, duplicated, fragmented or missing. For single-copy and duplicated classifications, the target gene must be present and >95% identical to the expected size and sequence of the reference in the database [60].

## Acknowledgements

We thank Paula Adams, Denise Akob, Louis Bubrig, Rebecca Varney and Kevin Kocot for helpful discussions. Patrick C. Phillips generously provided the PX356, PX439 and PX534 inbred strains and the Caenorhabditis Genetics Center provided the EM464 outbred strain. This work is funded by NSF uRoL 1921585 and NSF DEB 1941854.

## Author Contributions

Conceptualization, J.M.S. and J.L.F.; Methodology, J.M.S. and J.L.F.; Formal Analysis, J.M.S., J.D.M., A.C.M, and J.L.F.; Resources, J.L.F.; Writing – Original Draft, J.M.S., J.D.M. and J.L.F.; Writing – Review & Editing, J.M.S., J.D.M and J.L.F.; Visualization, J.L.F.; Supervision, J.L.F.

---

**SFigure 1.**
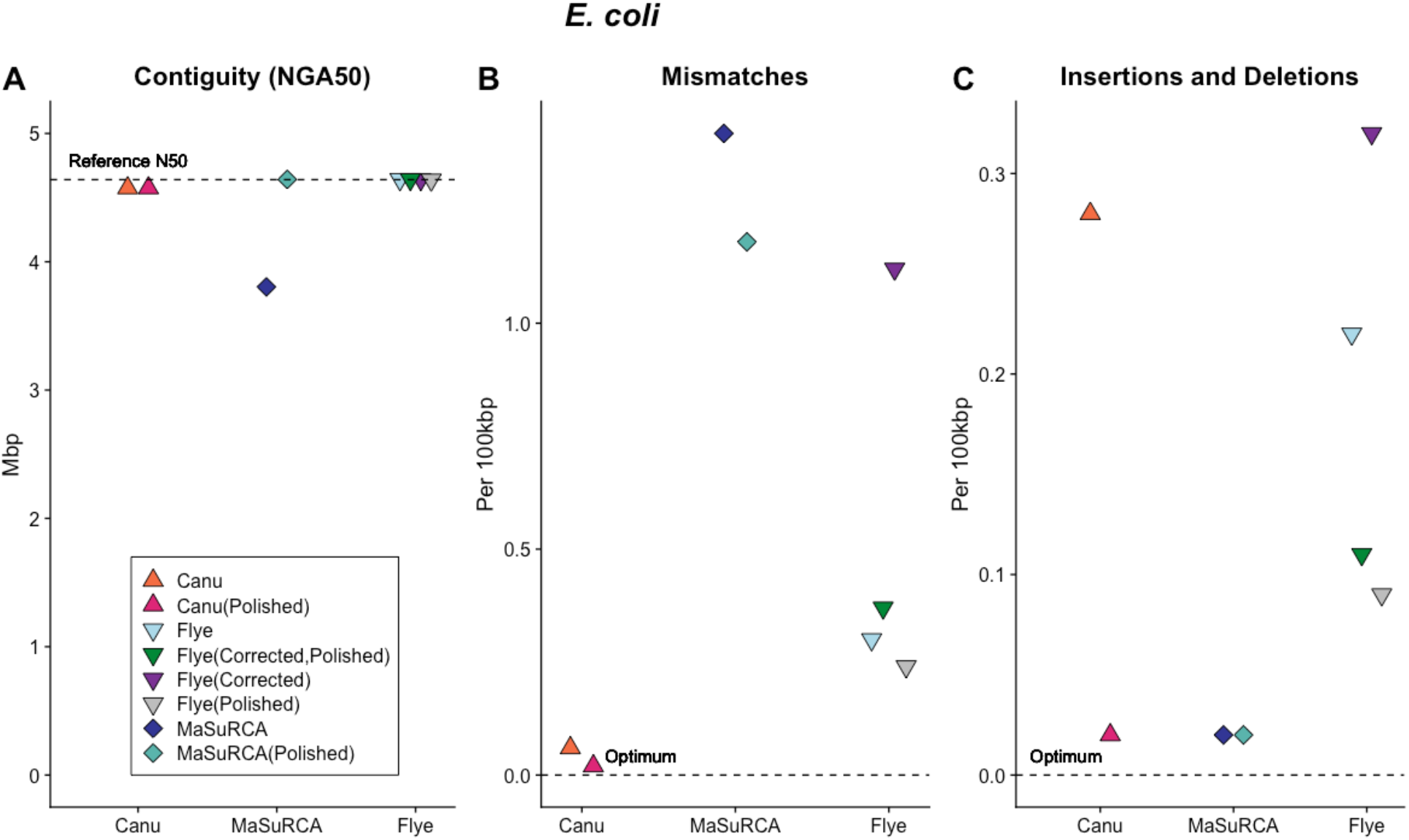
The relative performance of Canu, Flye and MaSuRCA assembly software with raw datasets, datasets with reads pre-corrected (‘Corrected’ in the figure legend) and post-assembly polishing with Illumina short reads (‘Polished’ in the figure legend) as measured by A) genome contiguity; B) mismatches and C) insertions and deletions relative to the *E. coli* reference sequence. Both Canu and Flye assembled highly contiguous sequences for all datasets and the MaSuRCA assembled sequence was highly contiguous post-polishing. The Canu assembled sequences had few mismatches and insertion/deletion errors were eliminated through polishing. The MaSuRCA assembled sequence had numerous mismatches that were not addressed through polishing and few insertions and deletions. The accuracy of all Flye assembled sequences was improved with polishing.

**SFigure 2.**
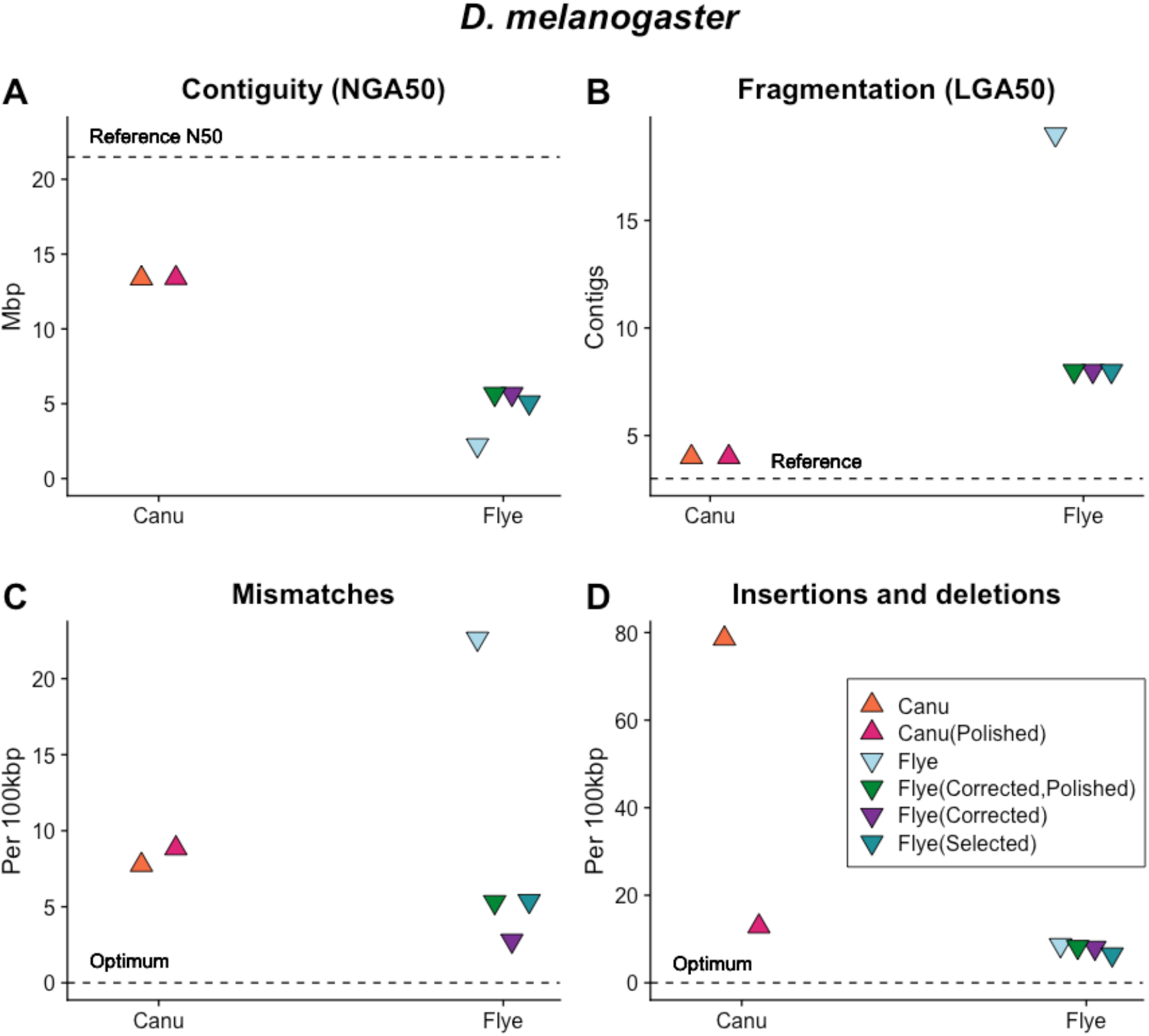
The relative performance of Canu and Flye assembly software with raw datasets, datasets with reads selected for length (‘Selected’), reads selected for length and pre-corrected (‘Corrected’ in the figure legend) and sequences post-assembly polished with Illumina short reads (‘Polished’ in the figure legend) as measured by A) genome contiguity; B) mismatches and C) insertions and deletions relative to the *D. melanogaster* reference sequence. Canu assembly of *D. melanogaster* ONT libraries produces assembled sequences with high contiguity, low fragmentation, few mismatches and relatively more insertions and deletions that are eliminated with post-assembly polishing. The performance of Flye can be increased by selecting, correcting and polishing.

**SFigure 3.**
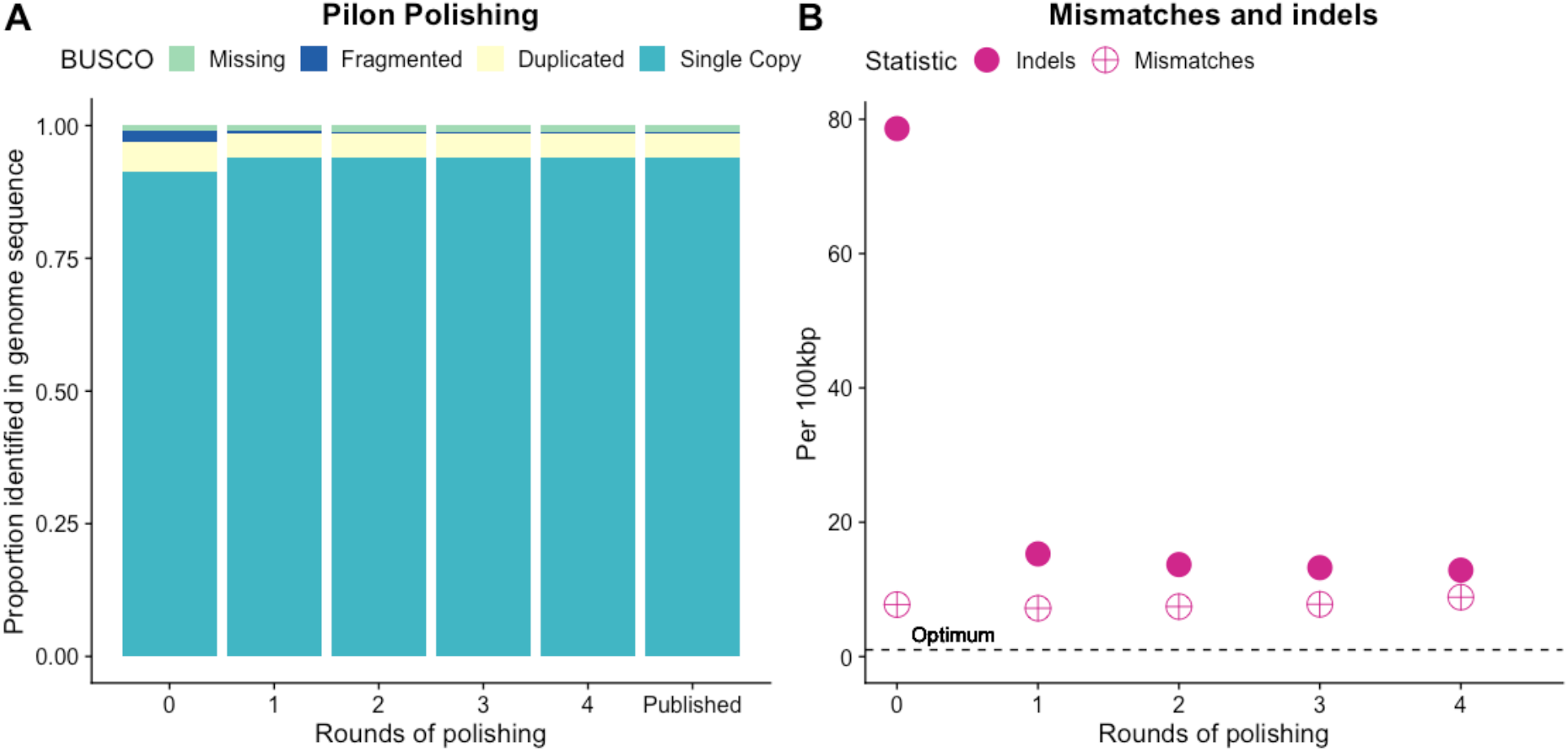
Polishing the assembled *D. melanogaster* sequence with Illumina libraries and the software package Pilon (REF) increases (A) the number of conserved genes found in single copy; and reduces (B) the number of mismatches and indels compared with the reference sequence. The bulk of this improvement occurs after 1-2 rounds of polishing.

**SFigure 4.**
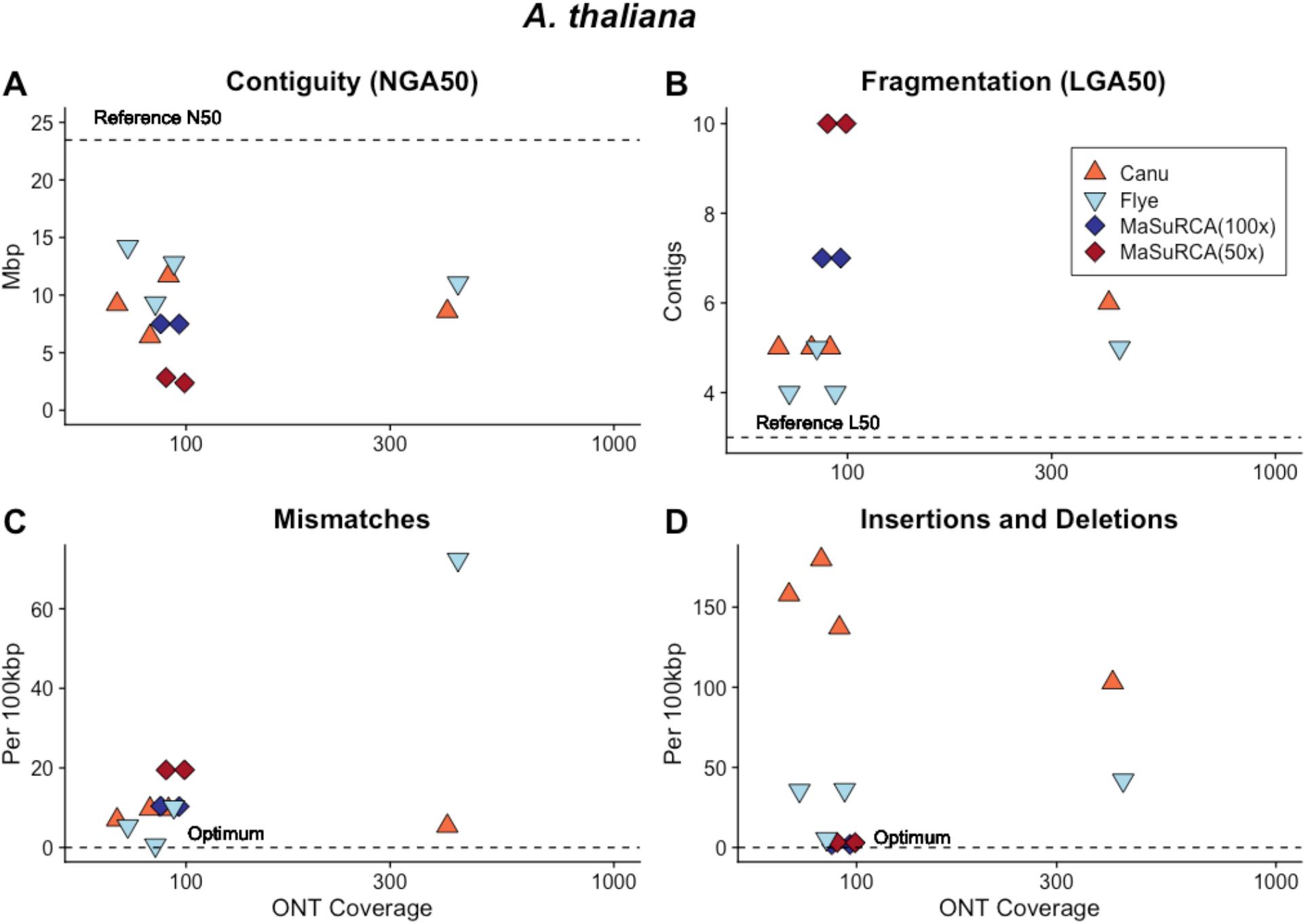
Canu, Flye and MasuRCA assembled sequences across a range of ONT sequencing coverages had A) low contiguity; B) high fragmentation; C) mismatches and D) insertions and deletions when compared with the *A. thaliana* reference sequence.

**SFigure 5.**
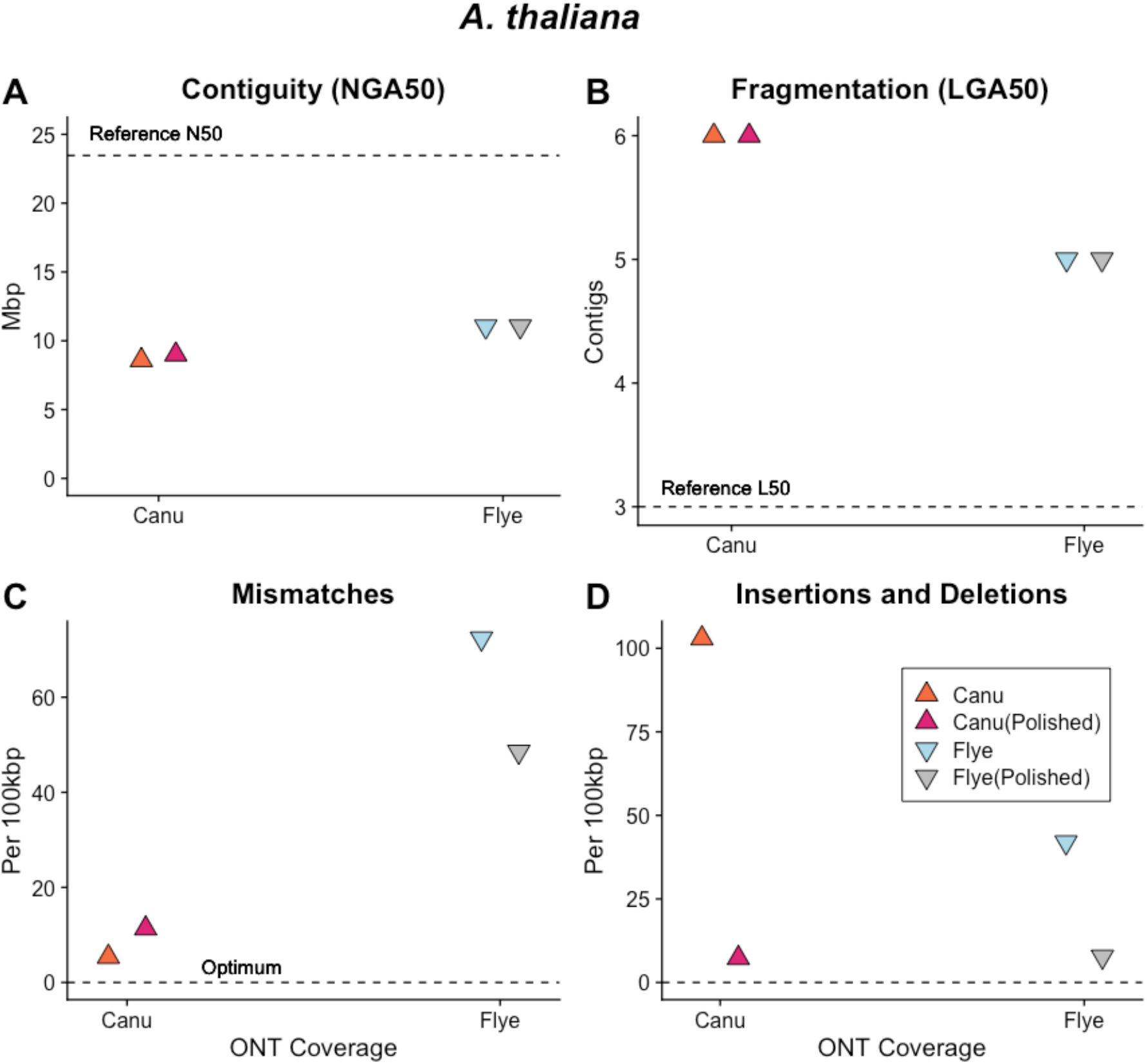
Post-assembly polishing increased the number of mismatches for Canu assembled sequences but decreased the number of insertions and deletions. Post-assembly polishing decreased mismatches, insertions and deletions for Flye assembled sequences.

**SFigure 6.**
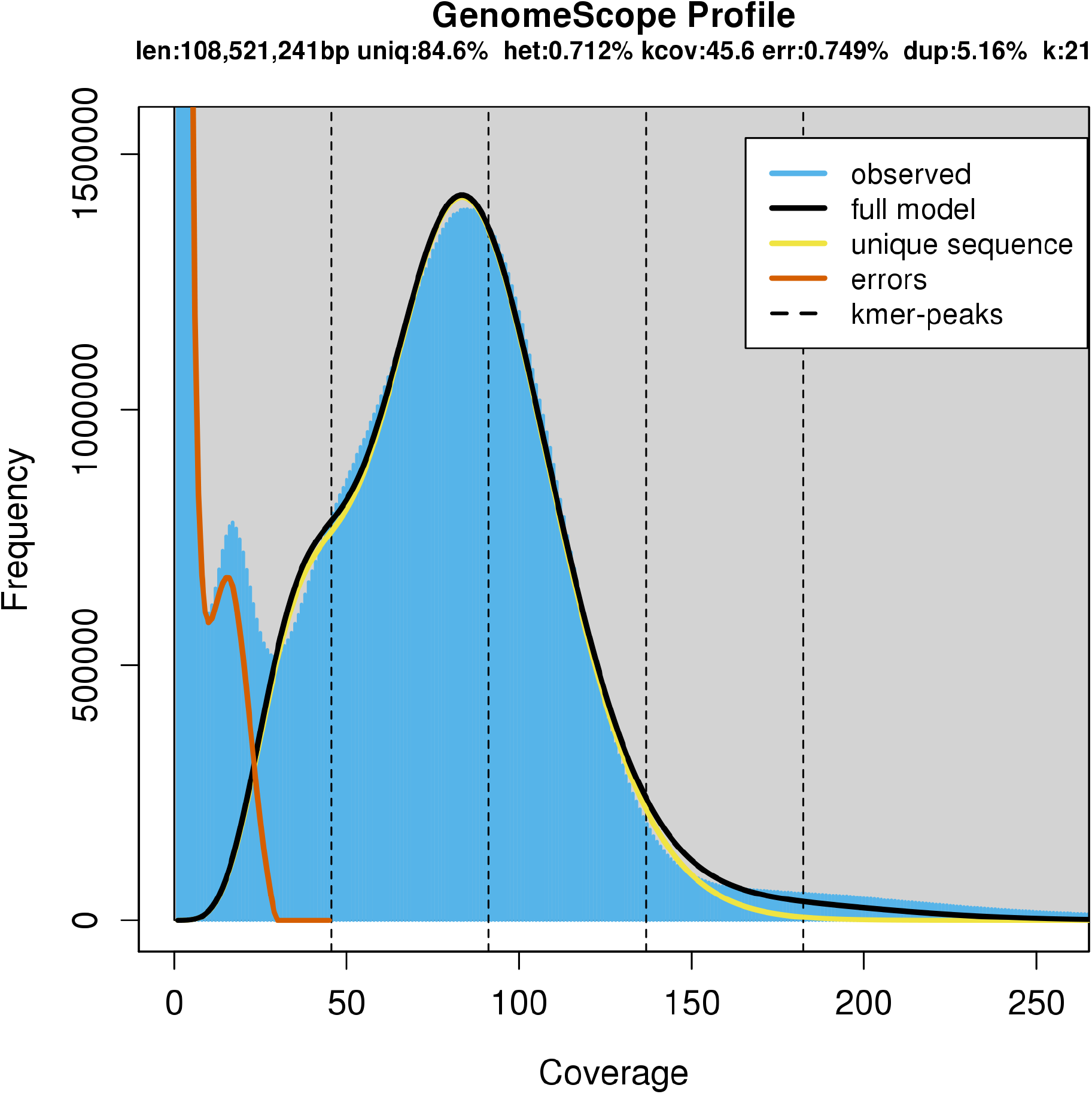
Analysis with the GenomeScope software revealed residual heterozygosity in the C. remanei PX356 sequences (estimated at 0.712%). Here, the residual heterozygosity is visible as a shoulder at ~50x sequencing coverage.

**SFigure 7.**
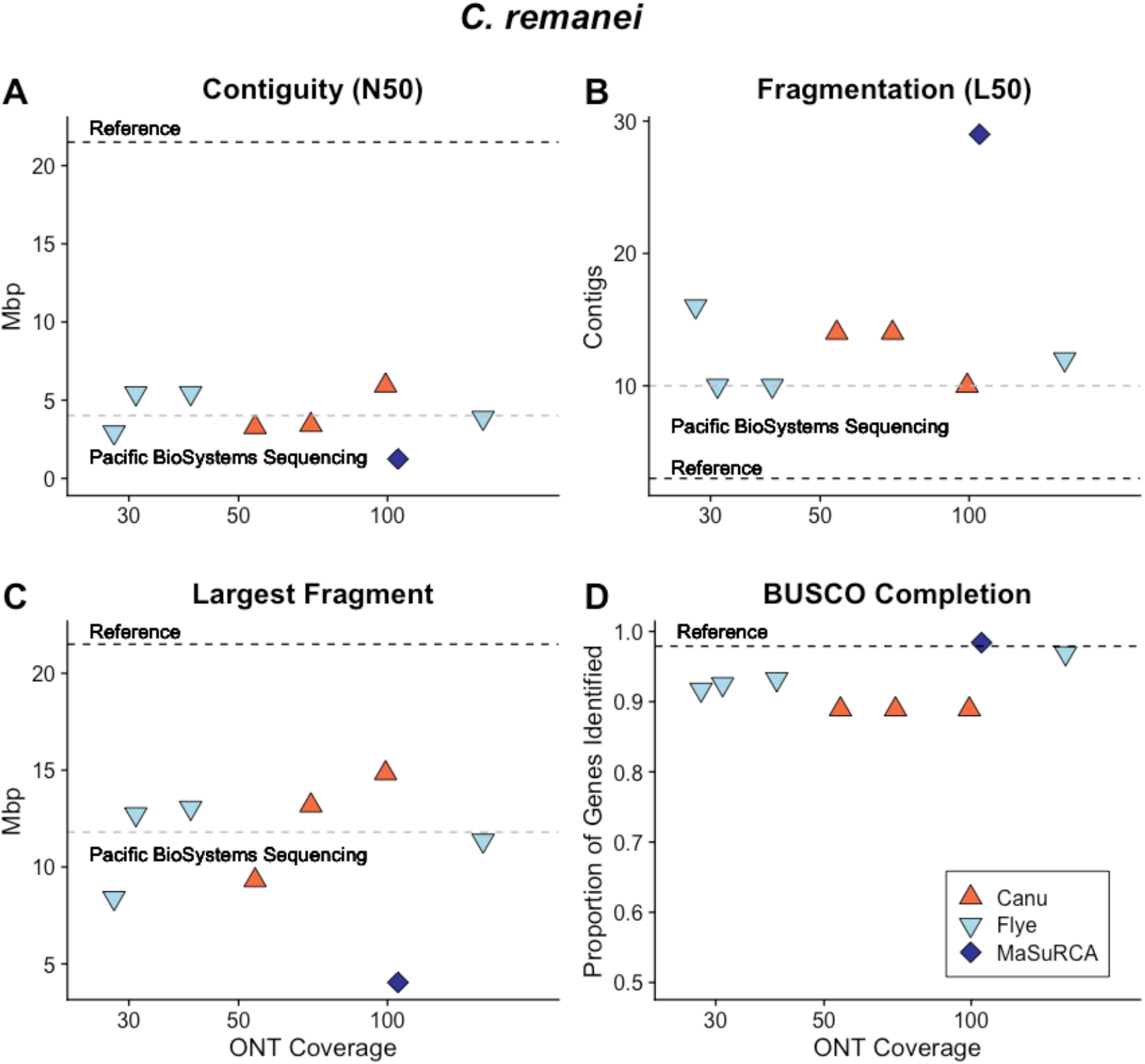
Canu and MaSuRCA assembled *C. remanei* sequences with A) low contiguity and B) high fragmentation compared with the true chromosome size and number but the sequences were of comparable contiguity and fragmentation when compared with Pacific BioSystems sequences. The C) largest fragment was also comparable to that achieved by Pacific BioSystems and the BUSCO genome completeness was high with assembly of a high coverage ONT library.

**SFigure 8.**
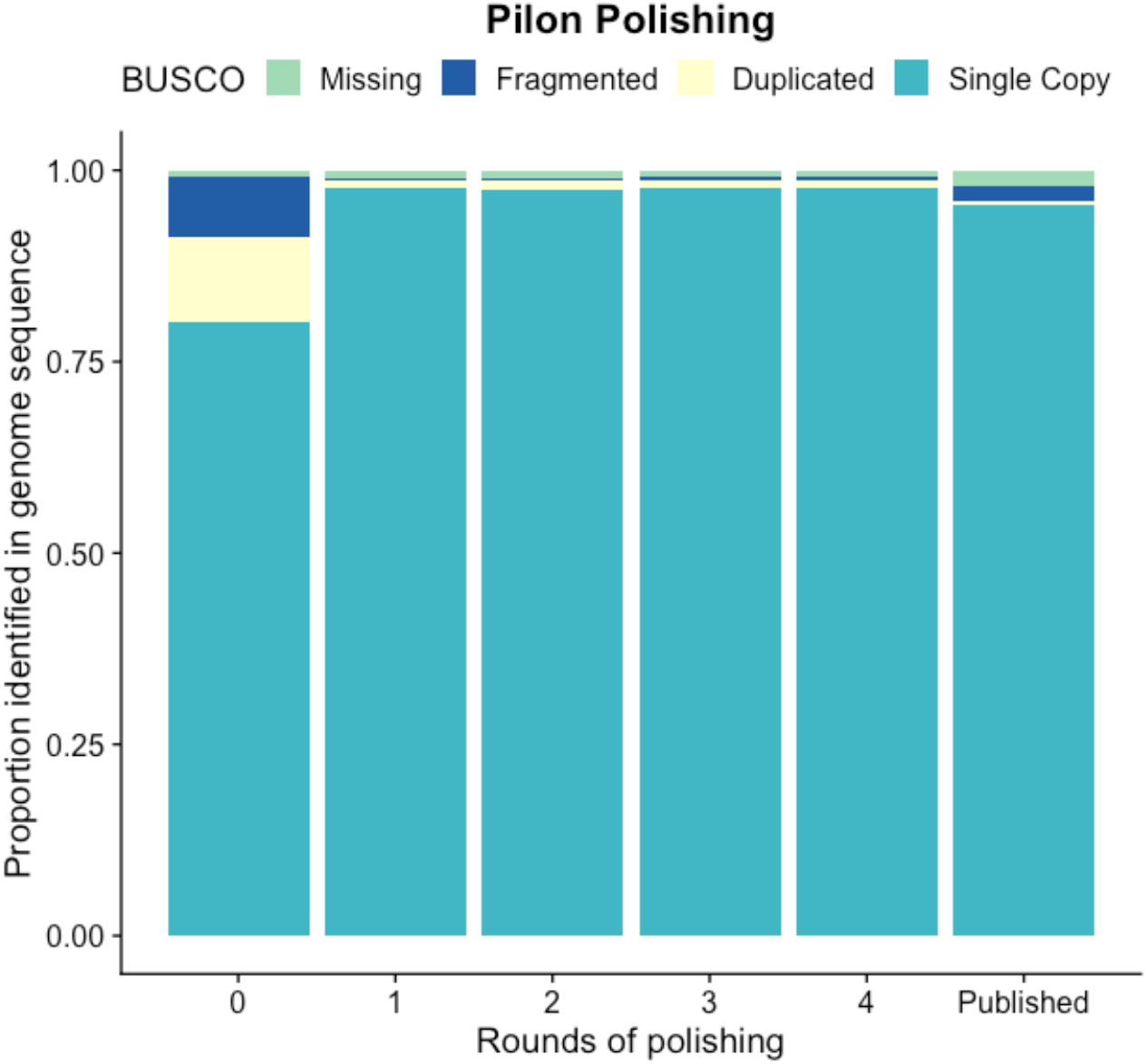
Polishing the *C. remanei* PX356 empirical assembled sequence with Illumina libraries and the Pilon software package (REF) increases the BUSCO completeness with a single round. The assembled sequence has a greater number of conserved genes found in single copy when compared with the published sequence (REF).

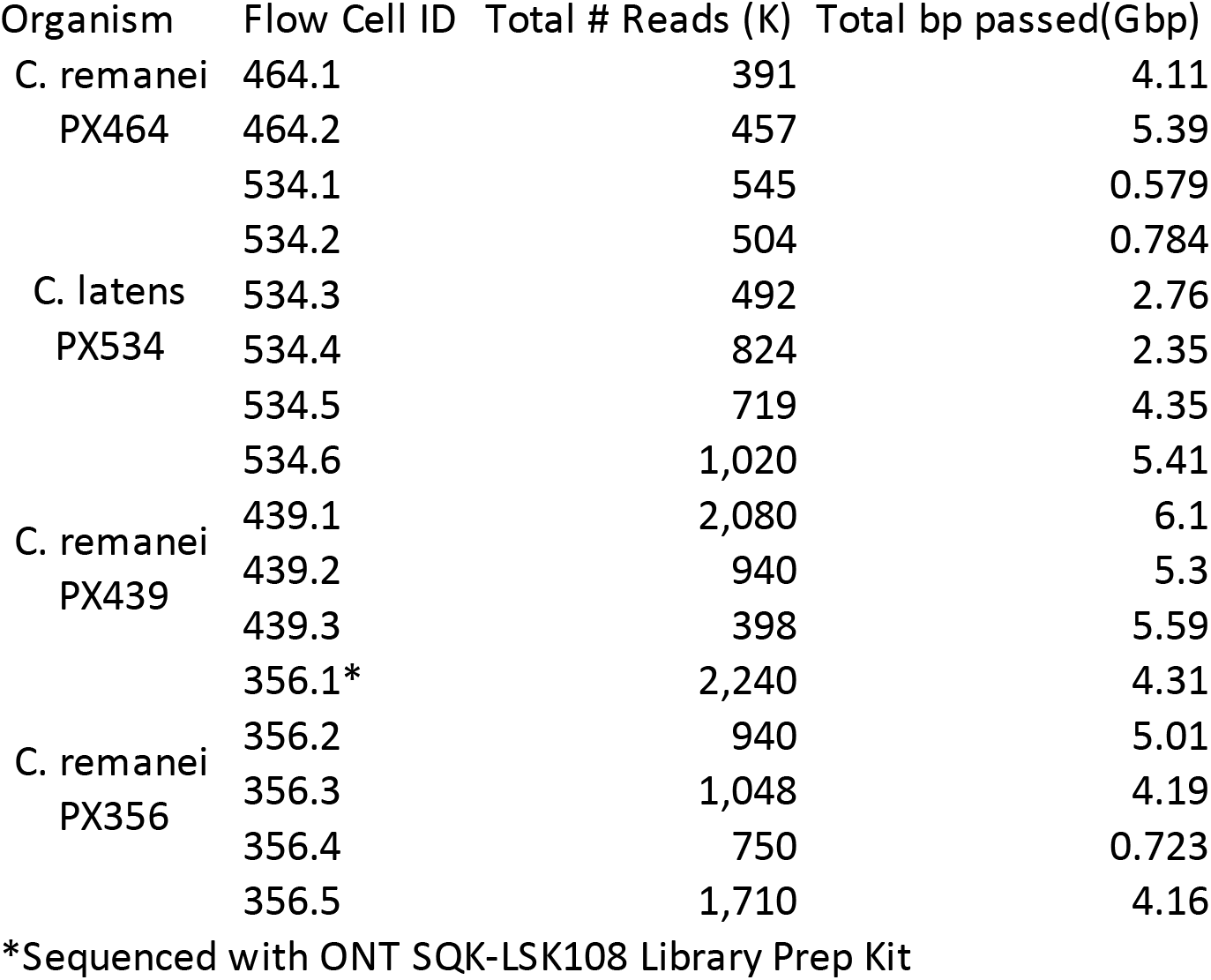

**STable 2.** Genome statistics for MaSuRCA-assembled Illumina paired sequences and error corrected Canu-assembled sequences. Although the Illumina paired end sequence is highly fragmented and incomplete it has fewer mismatches, insertions and deletions when compared with hybrid ONT and Illumina read sets.

| Sample | Read Coverage |  | BUSCO |  |  |  |  |  |
| --- | --- | --- | --- | --- | --- | --- | --- | --- |
|  | ONT | Illumina | Single (%) | Duplicated | Fragmented | Missing | Mismatches per 100Kpb | Indels per 100Kpb |
| *A. thaliana* | 0 | 100 | 98.1 | 0.9 | 0.5 | 0.5 | 10.99 | 1.03 |
|  | 420 | polished | 99.1 | 0.5 | 0 | 0.4 | 11.36 | 7.3 |
| *C. elegans* | 0 | 200 | 95.8 | 0.8 | 1.6 | 1.8 | 4.78 | 1.02 |
|  | 336 | polished | 97.7 | 0.6 | 0.4 | 1.3 | 0.73 | 0.88 |
| *D. melanogaster* | 0 | 100 | 93.5 | 1.5 | 0.4 | 4.6 | 10.01 | 1.7 |
|  | 108 | polished | 94 | 1.2 | 0.3 | 4.5 | 8.84 | 12.87 |
| *E. coli* | 0 | 100 | - | - | - | - | 1.7 | 0.04 |
|  | 62 | polished | - | - | - | - | 0.02 | 0.02 |

**STable 3:** Polishing the *C. remanei* assembled sequence with an Illumina library at 20x depth increases the percentage of conserved genes identified by BUSCO (Simão et al., 2018) after 3 rounds of polishing with Pilon (Walker et al., 2014). Polishing with a higher depth Illumina library (114x average sequencing coverage) produces similar results after 2 rounds of polishing with Pilon. The highest percentage of conserved genes found in single copy is achieved after 3 rounds of polishing with Pilon and a high depth Illumina library at 227x average coverage.

| Sample | Polishing |  | BUSCO |  |  |  |
| --- | --- | --- | --- | --- | --- | --- |
|  | Rounds | Coverage | Single (%) | Duplicated | Fragmented | Missing |
| **Canu** | 0 | 20x | 80.3 | 0.9 | 7.7 | 11.1 |
| **Assembled** | 1 | 20x | 97.2 | 1 | 0.5 | 1.3 |
| ***C. remanei*** | 2 | 20x | 97.3 | 1 | 0.4 | 1.3 |
|  | **3** | **20x** | **97.6** | **1** | **0.2** | **1.2** |
|  | 4 | 20x | 97.6 | 1 | 0.2 | 1.2 |
|  | 1 | 114x | 97.6 | 1.1 | 0.3 | 1 |
|  | **2** | **114x** | **97.6** | **1** | **0.3** | **1.1** |
|  | 3 | 114x | 97.5 | 1 | 0.3 | 1.2 |
|  | 4 | 114x | 97.5 | 1 | 0.3 | 1.2 |
|  | 1 | 227x | 97.6 | 1 | 0.3 | 1.1 |
|  | 2 | 227x | 97.5 | 1 | 0.3 | 1.1 |
|  | **3** | **227x** | **97.7** | **0.9** | **0.3** | **1.1** |
|  | 4 | 227x | 97.7 | 0.9 | 0.3 | 1.1 |

**STable 4:**
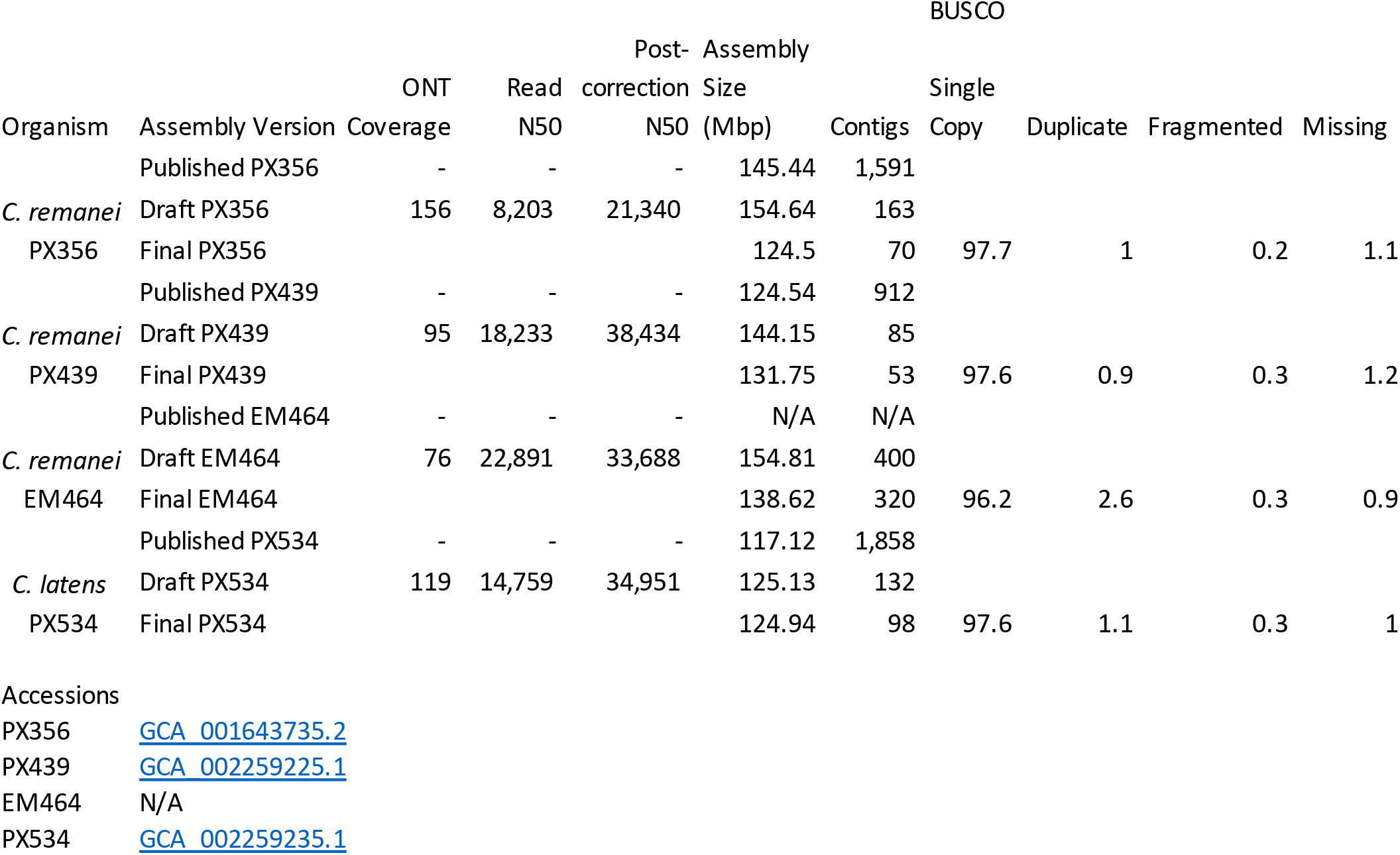

